# The Shape of Chemical Space

**DOI:** 10.1101/2025.09.23.678151

**Authors:** Krisztina Zsigmond, Akash Surendran, Lexin Chen, Ramón Alain Miranda-Quintana

## Abstract

The concept of chemical space is critical in cheminformatics, medicinal chemistry, and machine learning applications. Despite this, the high dimensionality of molecular representations greatly complicates its sampling, analysis, and visualization. A popular approach to overcome problem is to project these representations to a “human-manageable” subspace, usually containing only two dimensions. Non-linear dimensionality reduction techniques are by far the preferred strategy, following the reasoning that their flexibility can accommodate any arbitrary distribution originally present in the high-dimensional space. However, this ignores the elevated computational cost of these methods and the difficulty in tuning their hyper-parameters. Here, we show that basic properties of the metrics used in the original space can be used to infer the shape of the chemical space, which in turns suggests an optimal strategy to project chemical information to lower dimensions. The key insight is to realize that, no matter the set of molecules, their fingerprint representation can be considered to lie on a hyper-spherical surface. The smooth nature of this manifold means that we can use clustering to identify locally-dense sectors of chemical space, and selectively project them simply using linear (hyper-parameter free) methods, like principal component analysis. This approach surpasses non-linear techniques in several neighborhood preservation metrics, while only requiring a fraction of the computational cost. This pipeline is implemented in our N-Ary Mapping Interface (NAMI: https://github.com/mqcomplab/NAMI), which we tested in the visualization of 10 million molecules.

## 1. INTRODUCTION

It is a truth universally acknowledged, that a chemist in possession of a large dataset, must be in want of a non-linear projection method to analyze it.^1–7^ This is a consequence of the high dimensionality of the vector representations commonly employed to encode molecular information.^8–10^ Be it the thousands of bits in popular fingerprint formats or the often even larger arrays of molecular descriptors, it is not possible to directly interpret these vectors. This is particularly evident in the former case, since the similarity tools^11–14^ that are used to work with bit-vectors can be far removed from our intuition in handling metric spaces with Euclidean-like relations. This possess a key problem: how to represent the chemical space in a way that is easy to navigate and interpret? The ever-increasing size^15–20^ of synthetically and theoretically accessible chemical space complicates this task, since the computational cost of representation methods can quickly become prohibitive. Also, the effectiveness of ML-based models in chemistry is intrinsically linked to how these representations capture structure and property-based information.^21–24^ Molecules close together in the chemical space typically share similar properties or biological activities, which is critical for tasks like virtual screening and drug discovery.^25–34^ Ideally, ML techniques can learn these complex structure-property relationships by generalizing patterns across chemical space, enabling rapid prediction of properties like solubility, bioactivity, toxicity, etc., for untested compounds. Quantifying the distribution of molecules in chemical space can, in principle, help in identifying regions containing drug-like molecules. Hence, the challenge of chemical space exploration is to map out promising regions of this vast space to narrow down billions of possibilities into a manageable set.

Despite the aforementioned importance of the concept of chemical space in modern drug design and ML studies, it is worth noting that in the vast majority of cases, this notion is usually meant to simply be a “collection” of molecules, rather than an actual mathematical “space”.^35–39^ That is, virtually no pre-existing structure is considered when dealing with chemical libraries, which complicates their analysis. This is reinforced by the shortcomings found trying to devise more intuitive vector-inspired molecular representations. Going back to the problem of interpretability, if we want to visually inspect a compound dataset, we need to reduce it to a 2D or 3D set so the standard go-to dimensionality reduction (DR) methods used in chemical applications often ignore any pre-existing characteristics of the original high-dimensional space. As detailed below, the quality of the DR process is evaluated through the conservation of neighborhood information going from the high-to the low-dimensional representation. This is why methods like principal component analysis (PCA)^40–44^ are usually discouraged in favor of techniques like t-SNE,^45–50^ UMAP,^44,51–54^ and GTM,^55–58^ since the former can only do a linear projection, while the latter do not depend on any characteristics of the high-dimensional manifold. This, however, can come with a steep price: a well-known drawback of t-SNE and UMAP is that their flexibility demands a substantial computational cost. Moreover, an often ignored issue of these tools is that in their original formulations they assume a Euclidean relation between the original entries, and it is not trivial to accommodate the metrics used to study binary fingerprints. This has not damped the popularity of t-SNE and UMAP in chemical studies. For example, t-SNE has been used in visualizing the molecular and biological similarity of 2274 launched drugs from the Drug Repurposing Hub.^59^ Combining two t-SNE maps clustering these molecules based on molecular and chemical similarity, this method was used to predict the bioactivities of small molecules across a protein family. In another study,^48^ t-SNE was used to visualize reaction space, allowing chemists to visualize reaction types and explore synthetic pathways for drugs like commercial drugs like darunavir and montelukast. In a recent study, Orlov et al. explored the qualitative and quantitative efficiency of three non-linear dimensionality reduction methods: t-SNE, UMAP and GTM on chemical space visualization.^60^ A key issue associated with their methods is the hyperparameter selection problem. This issue arises due to the sensitivity of these algorithms to the set of hyperparameters, leading to significant impacts on the quality and interpretability of chemical space maps. There is no one-size-fits-all setting, since the optimal hyperparameters are highly dataset-specific due to varying molecular diversity, fingerprint type and dimensionality and sample size. Finally, exhaustive tuning is often infeasible for large datasets. For instance, t-SNE seeks to make pairwise similarities in high-dimensional and low-dimensional space as similar as possible by using the Kullback-Leibler (KL) divergence between these two distributions, making it an O(*N*^2^) process with the size of the training data. Recently, taking advantage of the hyperparameter dependence on the distribution of data, we showed that a small sampling (about 10%) of large datasets is enough to obtain optimal hyperparameters for these non-linear DR methods.^61^ Still, the problem of non-universality motivates the investigation of tools to extract maximum information about the shape of underlying chemical space, preferably with no hyperparameter tuning.

In this paper we explore two related issues: how simple properties of the metric used to analyze the high dimensional space can provide insights to improve the projections to lower dimensions, and how we can use clustering to make linear DR methods more accurate than t-SNE and UMAP. DR and clustering are usually presented together, especially for complex datasets. In fact, a common practice in recent times uses DR as a pre-processing step previous to performing the clustering. There are two key reasons behind this popular choice: First, by directly “seeing” the data in 2D it is easier for the user to tune the often complex clustering hyperparameters until an “adequate” partition is found. Second, reducing the number of dimensions of the data to be clustered reduces the overall computational cost. However, these arguments can (at best) masquerade deeper issues with both DR and clustering or (at worst) give outright erroneous results. The former can be seen in the issue of the computational cost. While it is undeniable that reducing the dimensions of the data representations will speed up the clustering, this is not the key factor preventing the scalability of clustering methods, since their cost is more heavily dependent on the number of molecules than in the dimensionality of their representation (not to mention, the latter is negligible compared to the former in most real-world applications). More importantly, the initial DR step can be detrimental by failing to capture the relations between the molecules in their original space, while biasing the user towards easy-to-spot patterns in low dimensions. In short: doing DR and then clustering is a great idea only if one already knows how the data should be partitioned. Here, we show that simply reversing this recipe (performing the clustering in the original space, and then doing the DR) considerably improves the quality of the final projections, to the point where PCA even surpasses non-linear alternatives. This not just allows to obtain better low-dimensional chemical representations, but it does so at a fraction of the computational cost of current approaches, to the point that multi-million libraries can now be fully processes in minutes in a single workstation. To conclude, we provide a package (N-Ary Mapping Interface, NAMI) that implements this pipeline, which is freely available at: https://github.com/mqcomplab/NAMI

## 2. METHODS & SYSTEMS

### 2.1 Local neighborhood metrics

As noted by Orlov et al.,^60^ DR studies need to be (carefully and quantitatively) evaluated on how well they preserve the information from the high dimensional space (basically, how good is a map that does not reflect the relations from the set to be mapped?). Here, we explore several indicators that quantify different aspects of this condition. Arguably the simplest of these criteria checks the consistency of the *k* nearest neighbors of each point in both these spaces, which is captured in the Average Nearest Neighbor Preservation (*P*_*NN*_). Another metric called Co-k Nearest Neighbor Size (*Q*_*NN*_) is used to track the shift in the rankings of the first *k* nearest neighbors during projection. The area under the curve (AUC) of the *Q*_*NN*_ curve then contains information about all possible neighborhoods and summarizes global information preservation. An even more detailed picture of local neighborhood structure is provided by Local Continuity Meta Criterion (LCMC), which is a normalized version of *Q*_*NN*_, favoring local information by penalizing distant neighbor calculations. Additionally, *Q*_*local*_ and *Q*_*global*_(respectively, the integral of the LCMC from 0 to a given *k*_*max*_, and from *k*_*max*_ to infinity) dissect the information contained in LCMC into local and global components. Finally, the quantification of false neighbors in the low dimensions can be done using two metrics: Trustworthiness and Continuity. Trustworthiness measures the proportion of points that are nearest neighbors in the projection but were not neighbors in the original space, penalizing false neighbors in low dimensions. Continuity, on the other hand, measures the opposite, points that were true neighbors but were not projected as such. The evaluation of these metrics was done with the help of code developed by the Varnek group, which is freely available at https://github.com/AxelRolov/cdr_bench.

### 2.2 Clustering large chemical spaces

The clustering algorithm to be used before the DR should ideally have the following properties: 1) do not assume any favored pre-distribution of molecules (we do not want to impose a fixed number of clusters on the set), 2) able to robustly identify tight subsets (the average similarity between molecules in a cluster should not be below the desired threshold), 3) computationally efficient (to facilitate processing large chemical spaces). For these reasons, we decided to focus on the recently proposed BitBIRCH.^62,63^ This method only takes one parameter from the user (a similarity threshold), and it is guaranteed to find clusters at least as compact as the given threshold. Regarding efficiency, BitBIRCH scales linearly with the number of molecules to be clustered, which makes it ideal to handle ultra-large libraries. BitBIRCH’s defining characteristic is the reduction of the information of the molecules contained in each cluster to a condensed bit feature (BF). For instance, for the *j* ^th^ cluster we do not need to store each of its fingerprints, instead, the BF must contain the number of molecules in the cluster, *N _j_*, the indices of the molecules, mols _*j*_, and a vector with the (linear) sum of the columns of the fingerprints belonging to the cluster, **ls** _*j*_. For simplicity, we also decided to store the cluster centroid, **c** _*j*_, although if preferred, this could be calculated on the fly. There are several ways to determine cluster membership in BitBIRCH (radial, through post-clustering refinement), but here, as a proof of principle, we focus on the diameter criterion. That is, a new molecule will be added to a cluster if the average of the Tanimoto similarities of the molecules in that set is above a user-specified threshold. Here, it is key to note that the average of Tanimoto values, ⟨T⟩, can be easily calculated with the information stored in the BF, through the iSIM framework:^64–67^

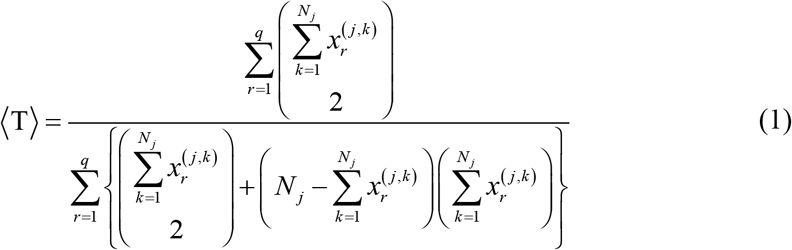

where 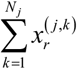 is the *r*^th^ component of **ls** _*j*_.

### 2.3 Cartography

As a simple model to study the projections of non-linear high-dimensional manifolds into planar surfaces (with particular relevance to fingerprint representations, as seen in section 3.1), we will consider the process of mapping the surface of the unit sphere in ℝ^*n*+1^ (centered at the origin of coordinates), defined as: *S*^*n*^ = {*v* ∈ ℝ^*n*+1^ : ⟨ *v*|*v* ⟩ = 1}. In particular, we will focus on the gnomonic and stereographic projection (Fig. 1).

**Figure 1.**
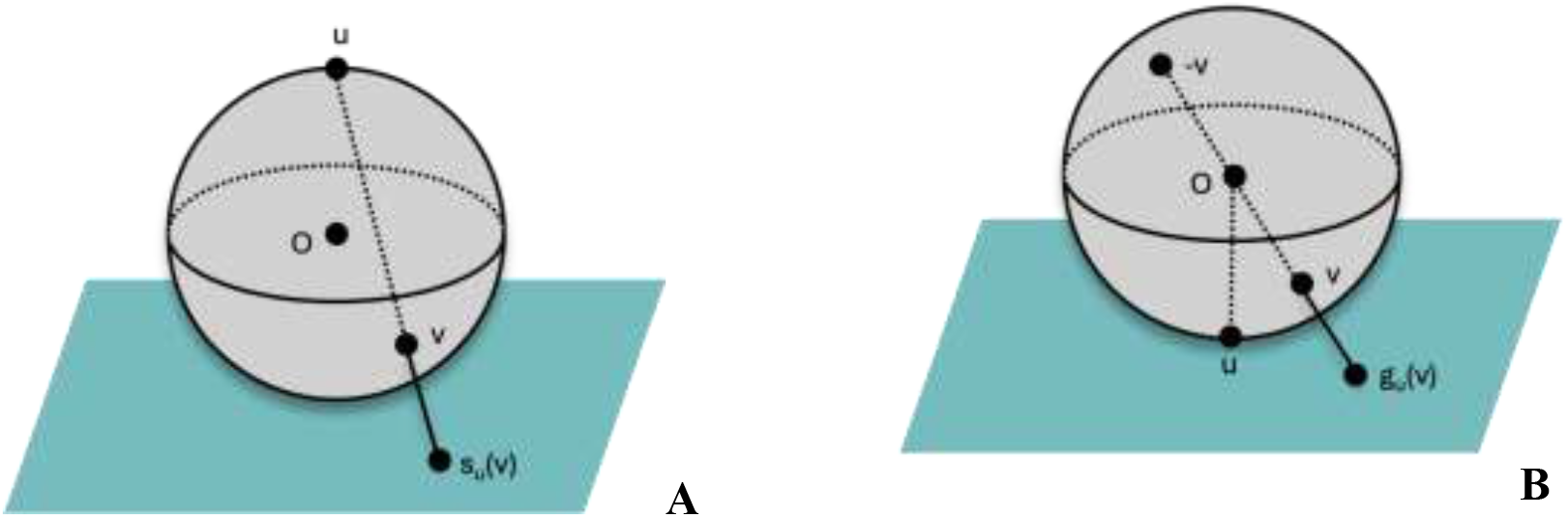
Cartoon representation of the stereographic (A) and gnomonic (B) projections.

The gnomonic projection is built starting from a vector *u* in *S*^*n*^ as:^68^

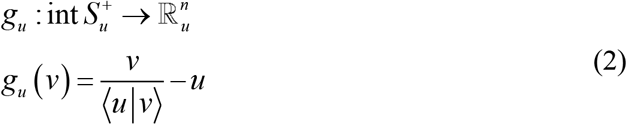

where 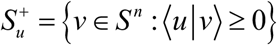.

Notice that although the projection is defined for vectors orthogonal to *u*, the projection of an orthogonal vector *v*, will have infinite components. (This happened for some of the molecules in the ChEMBL natural products set, when represented with ECFP fingerprints, see below.)

The stereographic projection^69^ is often defined with respect to one of the unit vectors (e.g., *e*_1_ = (1, 0,…, 0) ). In that case, the projection hyperplane ( ℝ^*n*^ ) is:

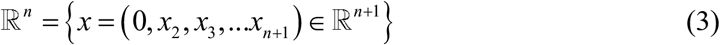

Resulting in the projection:

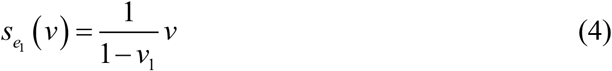

Where v1 is the component of v in e1. The stereographic projection then connects the points *u, v*, and *s*_*u*_(*v*) with a line (see Fig. 1A). Namely:

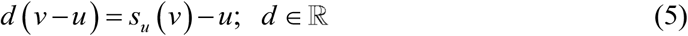

with:

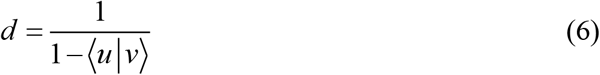

Here, we consider two possibilities for the reference vector *u*: the normalized (1, 1, 1, …) vector or the medoid of the set. Similarly, we use (-1, -1, …) and -medoid as references for the stereographic projection, since the projection plane in this case is on the opposite side of the unit sphere. Note that, to avoid over-complicating the ensuing discussions, we will still refer to these projections as stereographic 111 and stereographic medoid, instead of stereographic -1, -1, -1 and stereographic -medoid.

### 2.4 Datasets and DR methods

For the comparison of neighborhood preservation metrics across non-linear methods (t-SNE, UMAP and GTM) and PCA, a dataset CHEMBL219_Ki curated by Grisoni et al.^70^ on Dopamine D4 receptor containing 1859 molecules as SMILES strings was used. We used a small dataset to ensure that the comparisons could be completed within a suitable computational time, since non-linear methods like t-SNE and GTM do not scale linearly with number of molecules and hence could be infeasible for larger datasets.

The projections and local neighborhood analysis were performed on the CHEMBL33 natural products database with 64086 molecules.^71,72^ Two types of binary fingerprint representation were employed: Extended-Connectivity Fingerprints (ECFP)^73^ with radius 2 and 1024-bit length, and the 2048-bit RDKit fingerprints. Finally, to visualize a large dataset we use a random subset of ZINC20^74^ containing 10 million compounds, prepared primarily for Deep Docking Applications and provides clustered protonation ready SMILES strings. CHEMBL33 and CHEMBL219_Ki are available as csv files on the GitHub for this paper https://github.com/mqcomplab/NAMI and the ZINC20 dataset can be downloaded from the website: https://files.docking.org/zinc20-ML/smiles/ as ZINC_20_smiles_chunk2.

The hyperparameter grid for t-SNE, UMAP and GTM was chosen to be the same as Orlov et al. The t-SNE: perplexity: [1, 2, 4, 8, 16, 32, 64, 128] and exaggeration: [1, 2, 3, 4, 5, 6, 8, 16, 32], for UMAP: nearest neighbors: [2, 4, 6, 8, 16, 32, 64, 128, 256] and minimal distance: [0.0, 0.1, 0.2, 0.3, 0.4, 0.6, 0.8, 0.99] and for GTM: number of nodes: [225, 625, 1600], number of basis functions: [100, 400, 1225], regularization coefficients:[1, 10, 100] and basis widths: [0.1, 0.4, 0.8, 1.2]. The hyperparameters were tuned to preserve the maximum percentage of nearest 20 neighbors.

## 3. RESULTS

### 3.1 Tanimoto similarity and Euclidean distance

As noted above, one of the often-ignored problems plaguing DR studies is that even though the original fingerprints are compared using the Tanimoto index,^75^ the standard DR process itself starts by calculating the separation between compounds using the Euclidean distance. However, given that most data-processing algorithms (including DR methods) are optimized to operate over vectors with continuous entries, it is convenient to have a transformation that maps binary fingerprints to real vectors, while trying to preserve the ranking of similarities. To achieve this, we first have to consider the Tanimoto formula between two molecules:

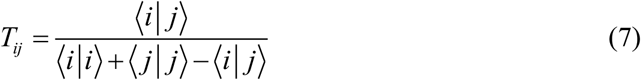

From here, it is clear that all molecules in “fingerprint space” are equidistant from a fingerprint consisting of only zeroes ( ∀*i*; ⟨0|*i*⟩ = 0 ). This provides a key revelation into the shape of chemical space, since the use of the Tanimoto similarity directly implies that the molecules lie in the surface of a multi-dimensional sphere. We can use this insight to take the binary representations to a proper metric space by simply normalizing the fingerprints. In other words, the transformation:

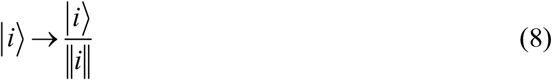

(where 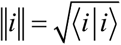) brings points in the surface of the “binary Tanimoto sphere” to the surface of the “continuous Euclidean sphere”.

Since ultimately our goal is to preserve the local environment as much as possible, we now ask the question: how capable is the *L*_2_ metric over the transformed (normalized) space of capturing the ranking of Tanimoto similarities? To check this, we looked at the relative monotonicity (e.g., consistency) of the Tanimoto and Euclidean values. In more detail: for multiple pairs of molecules, *i* and *j*, we calculate *E*_*ij*_(the Euclidean distance between molecules) and 1− *T*_*ij*_. If one of these values increases while the other also increases, both *T* and *E* rank the molecules in the same order. Also, to check the importance of Eq. (8), we computed the Euclidean distances over binary and normalized fingerprints.

Figure 2 shows the relative distribution between the Tanimoto and Euclidian values. As anticipated, the normalized fingerprints do a much better job at preserving the relations from the original binary space. Note the wide spread of values for both ECFP and RDKit fingerprints if the Euclidean distance is calculated with binary vectors. On the contrary, the distributions for the normalized vectors are much sharper, indicating a significantly better ranking preservation. As a way to quantify this, we calculated the Kendall correlation coefficients^76^ in all of these cases (see Table 1). Yet again, the utter inadequacy of the binary representation and the vastly superior performance of their normalized continuous counterparts is apparent for both types of fingerprints. It is remarkable that this level of agreement (over 0.9 Kendall correlation) is found just by enforcing perhaps the simplest condition in the chemical space, namely, going from the “Tanimoto sphere” to a “metric sphere”. This leads to the first key conclusion of this contribution: there is much to be gained by utilizing the underlying manifold structure of the chemical space. This has an immediate practical application: if we want to use analysis tools tailored for continuous representations, we do not need to code tailored versions that instead use Tanimoto similarity, we just need to normalize the fingerprints to ensure maximally-preserved relations between neighboring compounds.

**Table 1.** Kendall’s tau values between Tanimoto and Euclidean values for RDKit and ECFP fingerprints.

|  | RDKit | ECFP |
| --- | --- | --- |
| Binary fingerprints | 0.03 | 0.07 |
| Normalized vectors | 0.91 | 0.94 |

**Figure 2.**
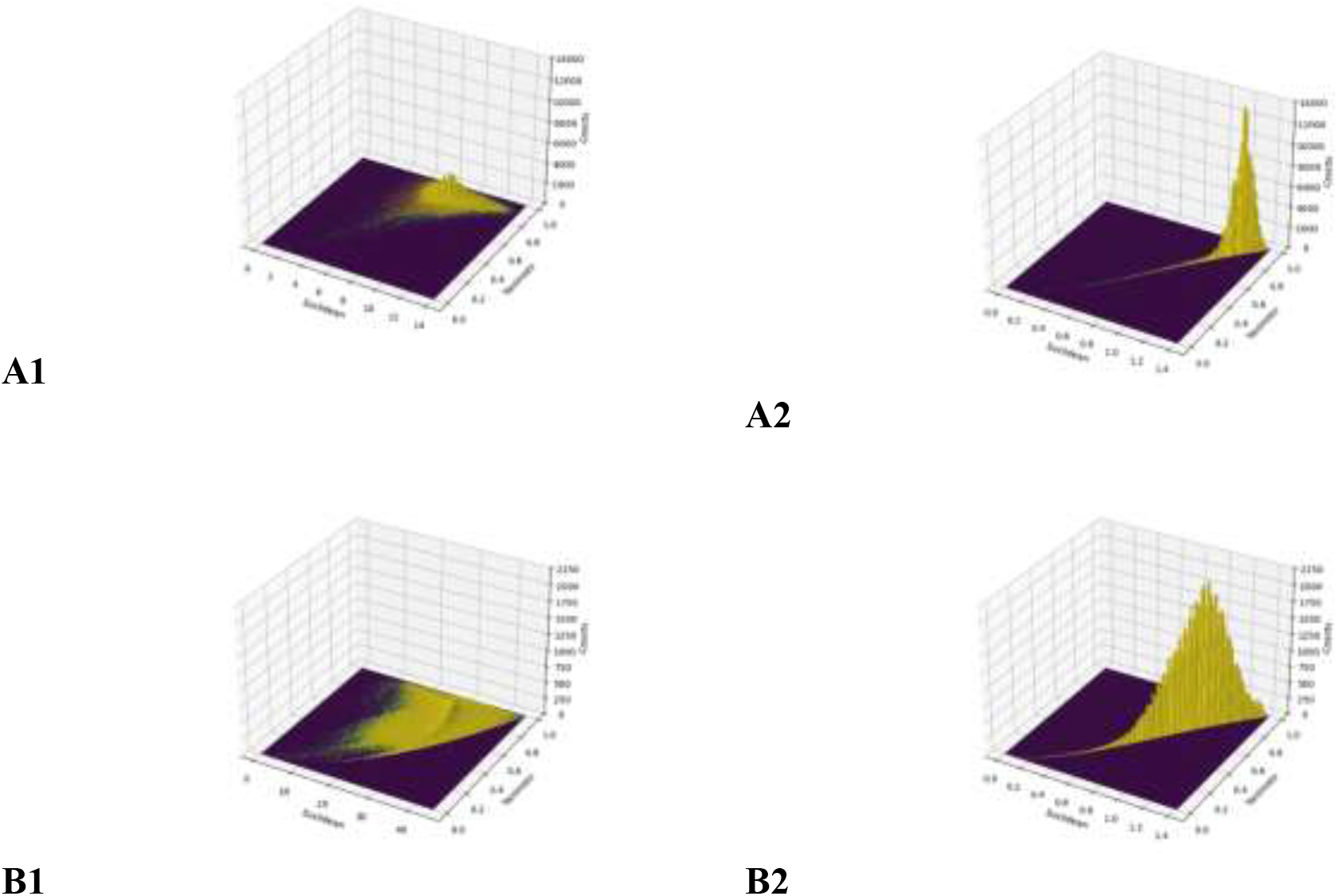
Distribution of Tanimoto and Euclidean values calculated for one million pairs of molecules. ECFP (A) and RDKit (B) fingerprints were considered, represented using binary (1) and continuous normalized (2) vectors.

### 3.2 Cartography and dimensionality reduction

The results from the previous section paint an enticing picture of a (hyper-)spherical chemical space with molecules resting in its surface. It is natural then to explore how this insight can be used to try to improve the DR results. In particular, we first focus on PCA. To this end, we tested three different protocols:

- **Naïve:** Performing the DR directly from the binary fingerprints.
- **Normalized:** Normalizing the fingerprints, and then performing the DR.
- **Projected:** Normalizing the fingerprints, projecting them into a hyper-plane, and then performing the DR.

In all cases, we chose to project to a two-dimensional space, since this is by far the most popular (and, arguably, convenient) way to visualize chemical space. The naïve method is included as a benchmark, since this is the current approach used in the community. The projected alternative is used to test is an intermediate modification of the chemical space manifold (from spherical to flat) before the DR results in any improvement.

Table 2 shows the scalar metrics for the information lost for the RDKit fingerprints with the 111 projections (the SI has the results for the medoid-centered projections and for the ECFP fingerprints). First of all, at the end of the Table, we once again corroborate the results presented in the previous section about the great performance of the normalized vectors (before reducing the dimensions to two). Note how just the normalization, or the normalization and then the gnomonic or stereographic projections show markedly high values for all the studied indicators. Notably, while both hyperplane projection methods show essentially identical results for the RDKit fingerprints, the stereographic projection is significantly better than the gnomonic for ECFP fingerprints (see the SI section). As expected, however, the distortion caused by the transformation from a spherical to a flat surface, carries an inescapable decrease in quality of the preservation of the local environments, as we can see by comparing the pure stereographic and gnomonic results with the metrics after performing just the normalization.

**Table 2.**
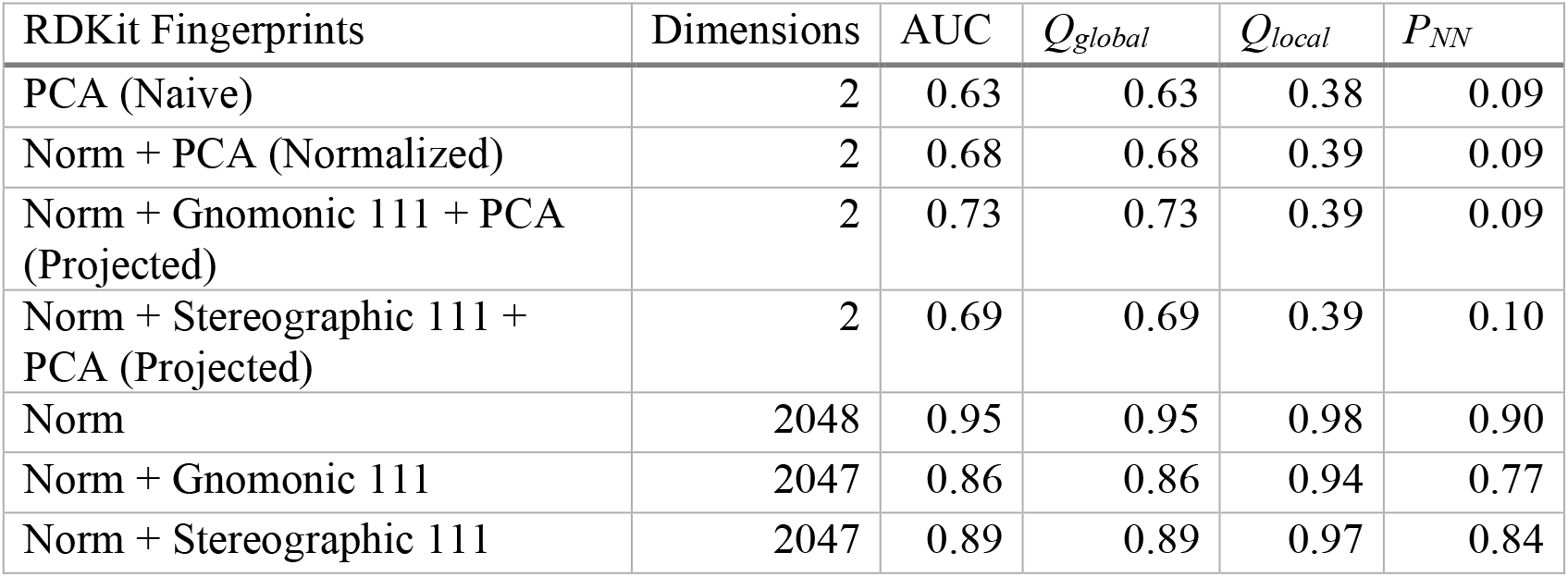
Local and global metrics for different combinations of hyper-plane projections, normalization, and PCA.

The results at higher dimensions provide further insights to analyze what happens when we perform PCA to go down to a bi-dimensional representation. Notice how normalizing before PCA provides a slight improvement for all of the considered metrics, as opposed to just doing PCA on top of the original binary fingerprints. Now, however, there is a difference between the hyperplane projections, with the gnomonic results being slightly better. These results are certainly encouraging, but unfortunately they are not as marked for the ECFP fingerprints (SI), where the normalization results in 2D are virtually equivalent to those obtained with the original fingerprints. Similar results are observed when the projections are centered around the medoid of the set (see SI).

Figure 3 shows the results for the other neighborhood preservation indicators. Here, it is easier to see the impact of the normalization. For instance, for the RDKit fingerprints, PCA after normalization provides better results than just doing PCA without normalizing for all descriptors and hyper-plane constructions. This is also the case for ECFP fingerprints (see SI), with the exception of the continuity index, which underperforms after normalizing the fingerprints.

**Figure 3.**
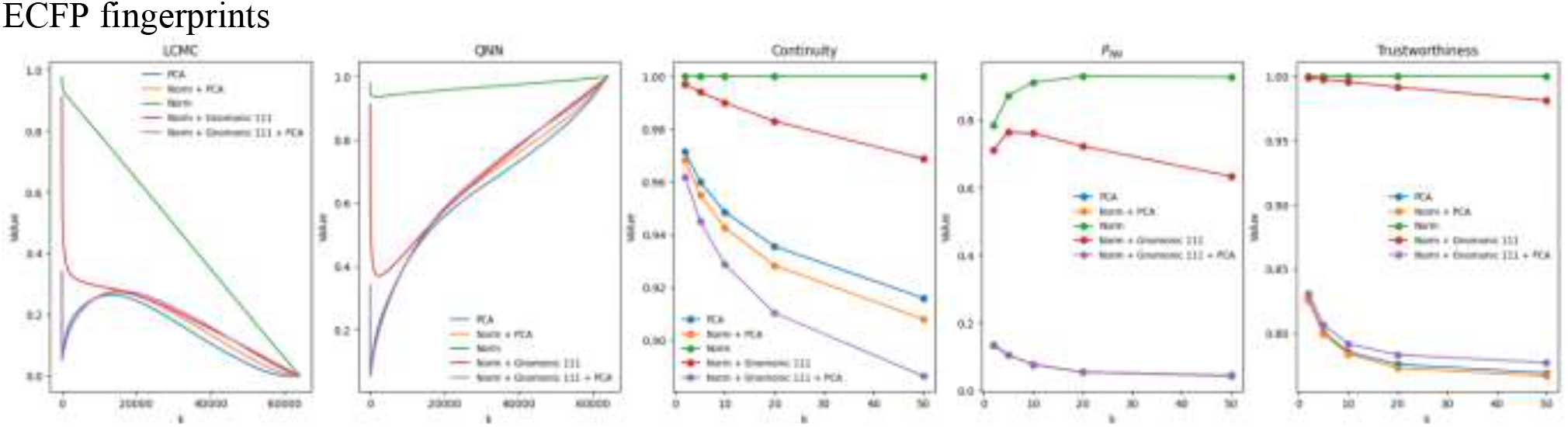
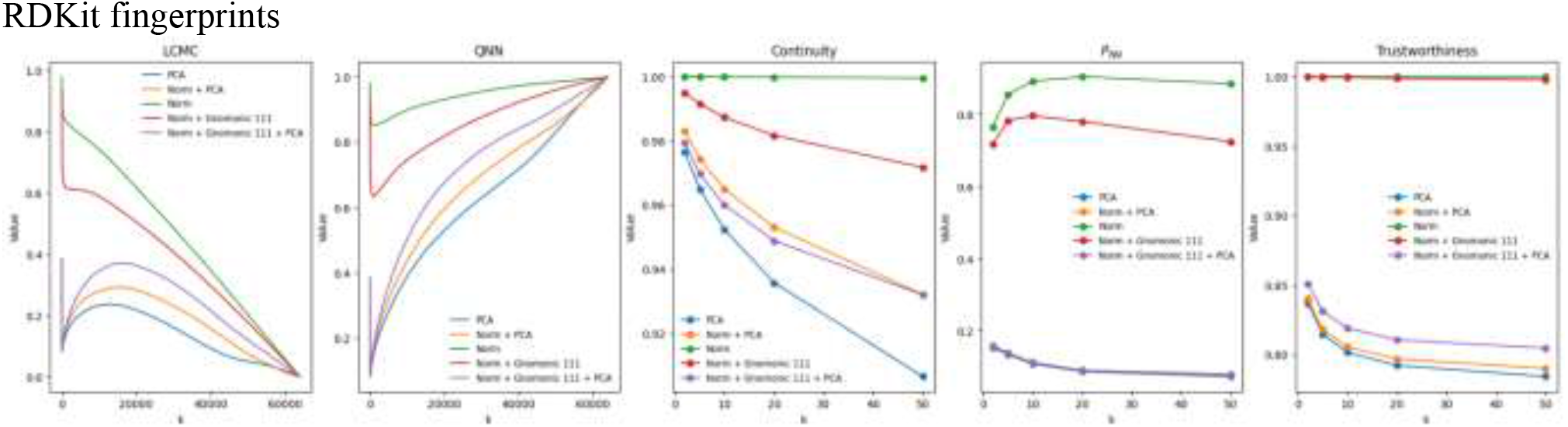
Neighborhood indicators for RDKit and ECFP fingerprints for the gnomonic projection.

Up to this point, we have focused on the impact of the normalization and hyper-plane projection for PCA, a linear DR method. It is of course of interest to evaluate the impact of these techniques on non-linear algorithms, like t-SNE, UMAP, and GTM. Considering the time constraint associated with non-linear DR methods, we performed this analysis on a smaller dataset containing only 1859 molecules. Still, in this very small dataset, we already observed noticeable differences in the time it took to process it with the different methods: PCA naturally consumed the least time to project the data, with just 6.5 seconds, while UMAP took about 3 minutes to do the same, and t-SNE and GTM required 19 minutes and 1 hour 10 minutes respectively.

Figures 4-5 compare some of the neighborhood metrics of the ECFP fingerprints of this dataset over the set of hyperparameters discussed in Section 2.4, which serve to illustrate some key points. First, the spread in the violin plots shows the distribution of data across different sets of hyperparameters, with a larger spread indicating a diverse range of values for the given metric. One can immediately observe that the global metrics (Fig. 4) are much more stable to variation in hyperparameters in comparison to the local neighborhood preservation metrics. For AUC (Fig. 4A), the average performance is consistent across all DR methods over all types of projections and shows similar values on average. All the non-linear methods just marginally outperform PCA in their best settings. GTM is the most stable across the hyperparameter set, showing a slight deviation (less than 0.04) across the range, while the performance of t-SNE and UMAP shows larger variations. Similar consistency in values can be observed for the case of *Q*_*global*_ as well (Fig. 4B) AUC and *Q*_*global*_ show similar violin shapes indicating a small *k* value for the inflection point (*k*_*max*_) of LCMC curve, hinting at a small *Q*_*local*_ contribution to the *Q*_*NN*_ curve. Overall, all these methods succeed in preserving more than 50% of global information, independent of the set of hyperparameters, hence providing a convincing picture of the global shape of the underlying chemical space.

**Figure 4.**
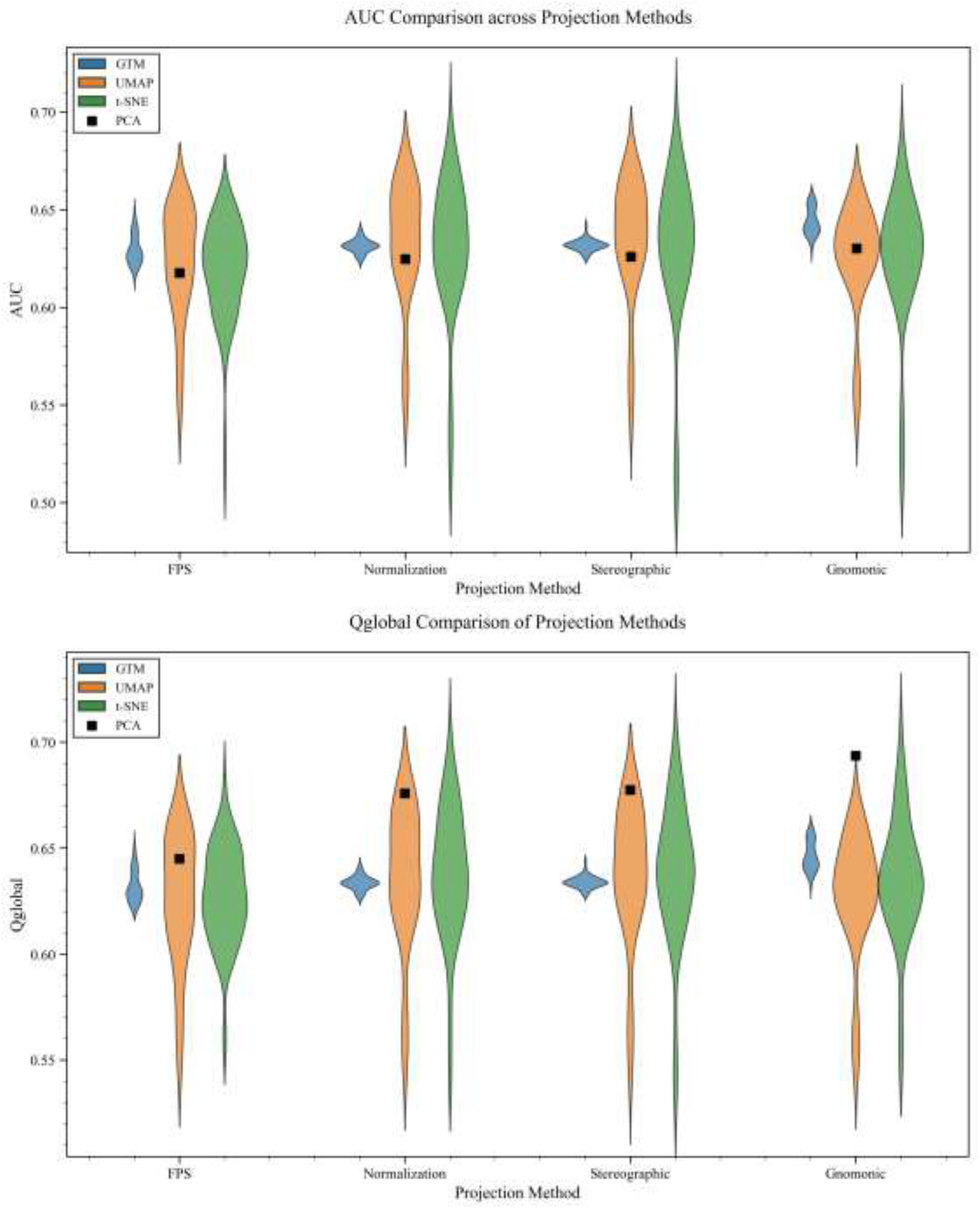
Global metrics for GTM (blue), UMAP (orange), t-SNE (green), and PCA (black) for CHEMBL219_Ki with ECFP4 fingerprints.

**Figure 5.**
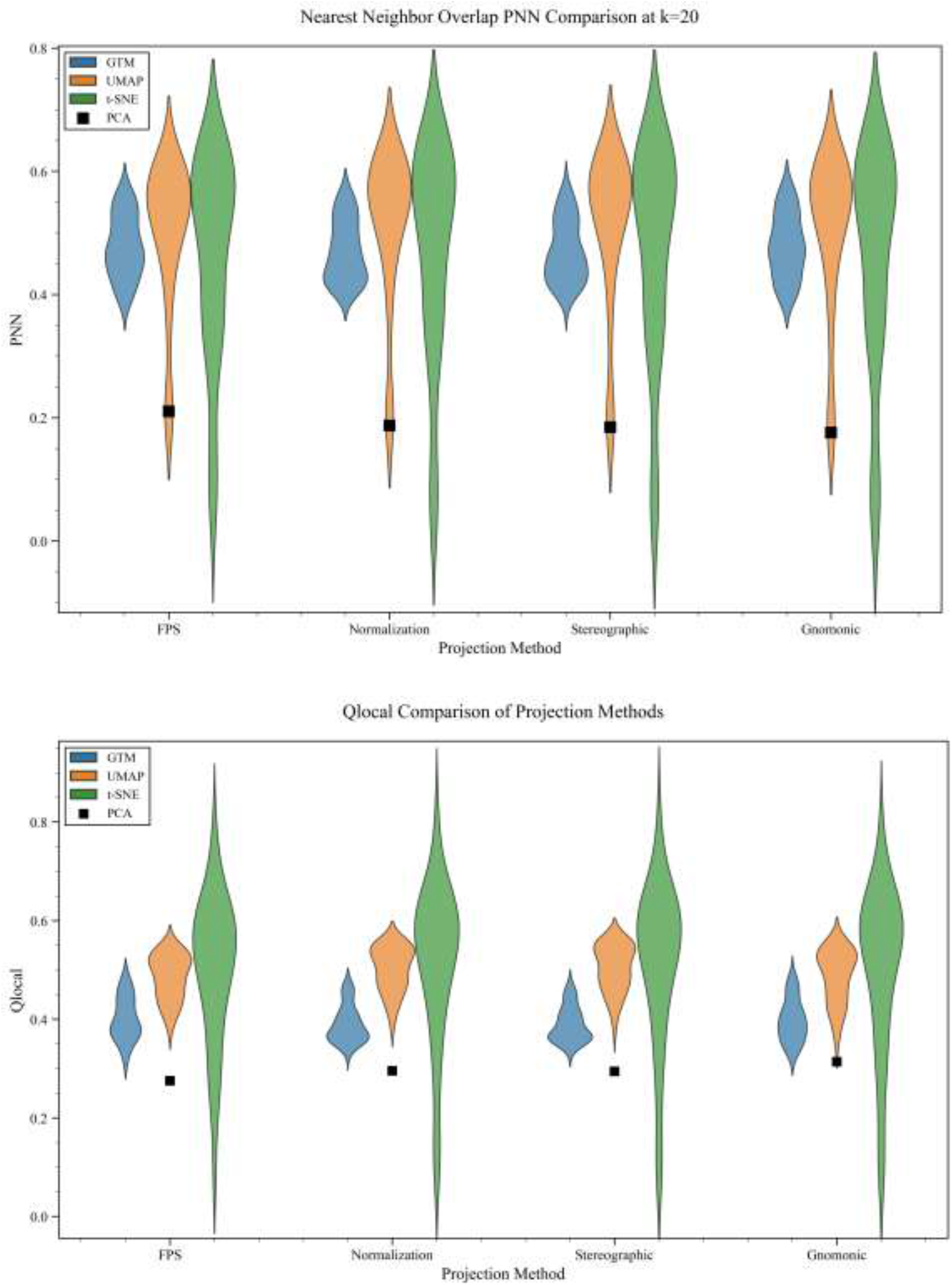
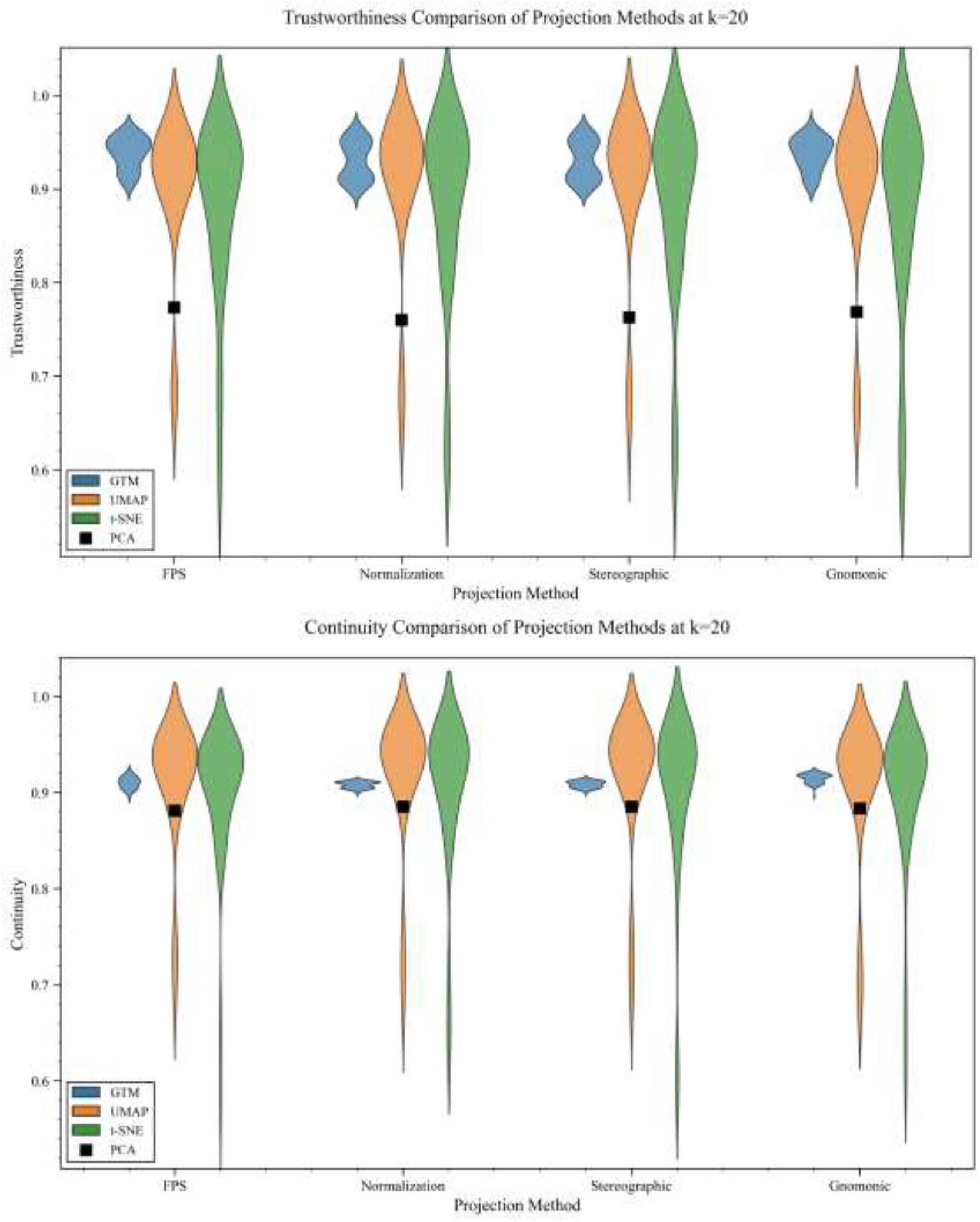
Local metrics for GTM (blue), UMAP (orange), t-SNE (green), and PCA (black) for CHEMBL219_Ki with ECFP4 fingerprints.

Local neighborhood preservation metrics, on the other hand, show a different picture (Fig. 5). The plot of *Q*_*local*_ (Fig. 5A) reveals that even though non-linear methods are still very sensitive to the choice of initial parameters, they significantly outperform PCA. In particular, t-SNE presents an interesting duality: it is both the best and worst method, with a 70% difference between the optimal and worst hyperparameter set, respectively. Similar results can be observed for the trustworthiness (Fig. 5B) and continuity (Fig. 5C) criteria. Yet again, PCA severely underperforms compared to the non-linear methods, which in turn can provide very good results, if their hyperparameters are carefully chosen. There is, however, another interesting characteristic of the non-linear methods, present both in the global and local metrics, and that is their consistency across the hyperplane projections that we saw could actually improve the PCA results. In other words, neither the normalization nor the hyperplane projections seem to improve the performance of t-SNE, UMAP, or GTM. A plausible explanation for this is that these methods are already extremely flexible, so considering some underlying structure in the high-dimensional fingerprint space does not appreciably impact their outcome. This leads to an obvious question: is it possible to use the insights from the “spherical” distribution of molecules in the high-dimensional space to develop a linear method that could compete with the non-linear projections? The following section responds this in the affirmative.

### 3.3 Project-after-clustering

The realization that the original fingerprint space endowed with a Tanimoto metric can be put in almost perfect correspondence with a spherical surface with Euclidean distances provides a nice insight into the structure of chemical space, but as noted above, it does not immediately lead to a dramatic improvement in the quality of the projected results. This is easy to rationalize if we remember that the much simpler problem of projecting the surface of our 3D planet into a 2D plane inescapably leads to very distorted regions. Essentially, no matter the nature of the starting (non-linear) manifold, any DR technique will inevitably lose and/or distort information, which is even more pronounced when we go from thousands of dimensions (fingerprints) to just two. (Not in vain, isometric embedding theorems from non-linear manifolds always require *increasing* the number of dimensions.) However, the potential solution to this issue can be found in how we approach mapping the surface of our planet: while it is possible to have approximate representations of the whole planet, whenever we need extra precision to navigate a given city, we only focus on that portion of the spherical surface, and only project that terrain. Essentially, the smaller the spherical portion that we want to map, the better the approximation of just using a linear (tangential) plane to that section. To apply this insight to chemical spaces, however, we need to find a way to first identify dense (e.g. “city-like”) regions in fingerprint space. This calls for a paradigm shift in the relation between clustering and DR: traditionally, DR is performed before the clustering, as a way to reduce the computational cost of the latter. Now, we propose doing the clustering first, as a way to improve the quality and preserve the information of the DR. This project-after-clustering (PAC) recipe also has the advantage of not requiring any hyperparameter tuning, beyond the selection of the neighborhood size for the projection. Of course, this paradigm could be coupled to any DR technique, but it is certainly more attractive to focus on simpler, linear, methods.

In order to establish a baseline to gauge the effects of the clustering approach, in Fig. 6 we “zoom” into the results of PCA and UMAP (done in the traditional way) for the 64K set (we only compare to UMAP because the other non-linear techniques took too long to converge for this library). Yet again, AUC, *Q*_*global*_, and *Q*_*local*_ show relatively similar performances for both DR strategies. On the other hand, *P*_*NN*_ and trustworthiness show a marked improvement for UMAP (when we use the best possible hyperparameters).

**Figure 6.**
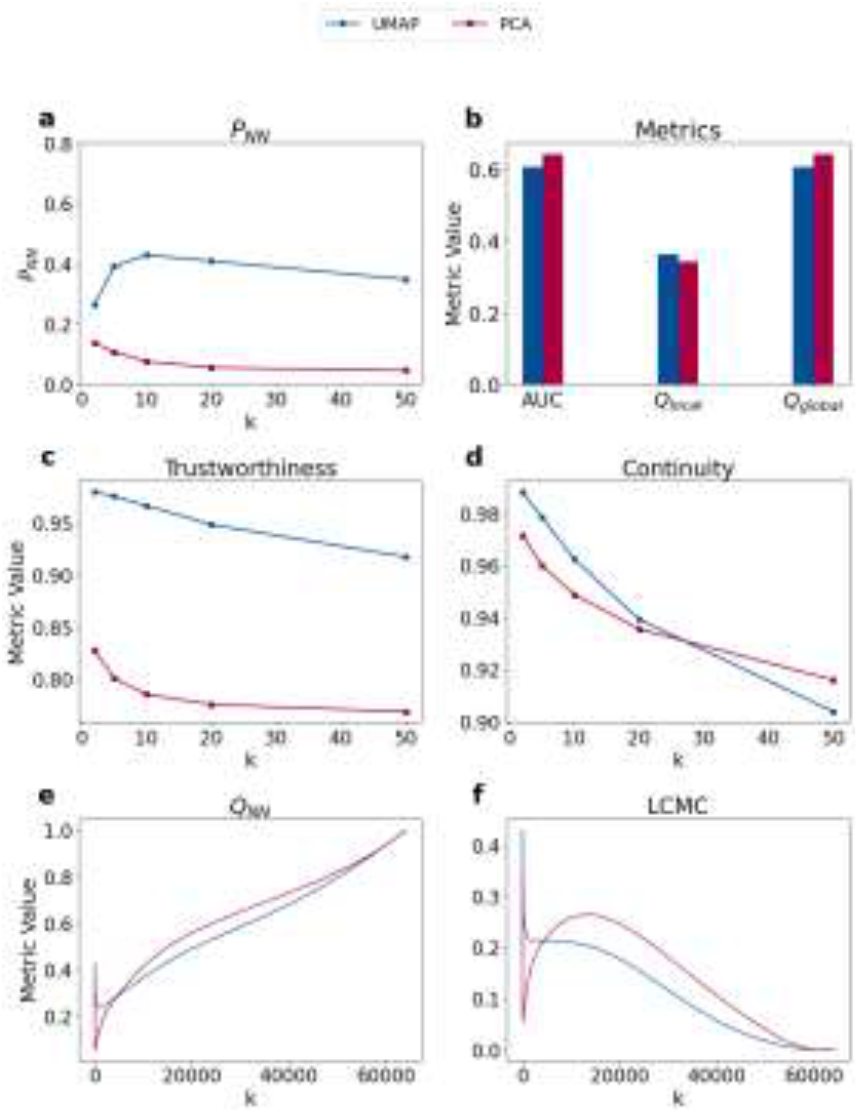
Detailed analysis of neighborhood metrics for the 64K set after projecting using PCA and UMAP for CHEMBL33 with ECFP4 fingerprints.

**Figure 7.**
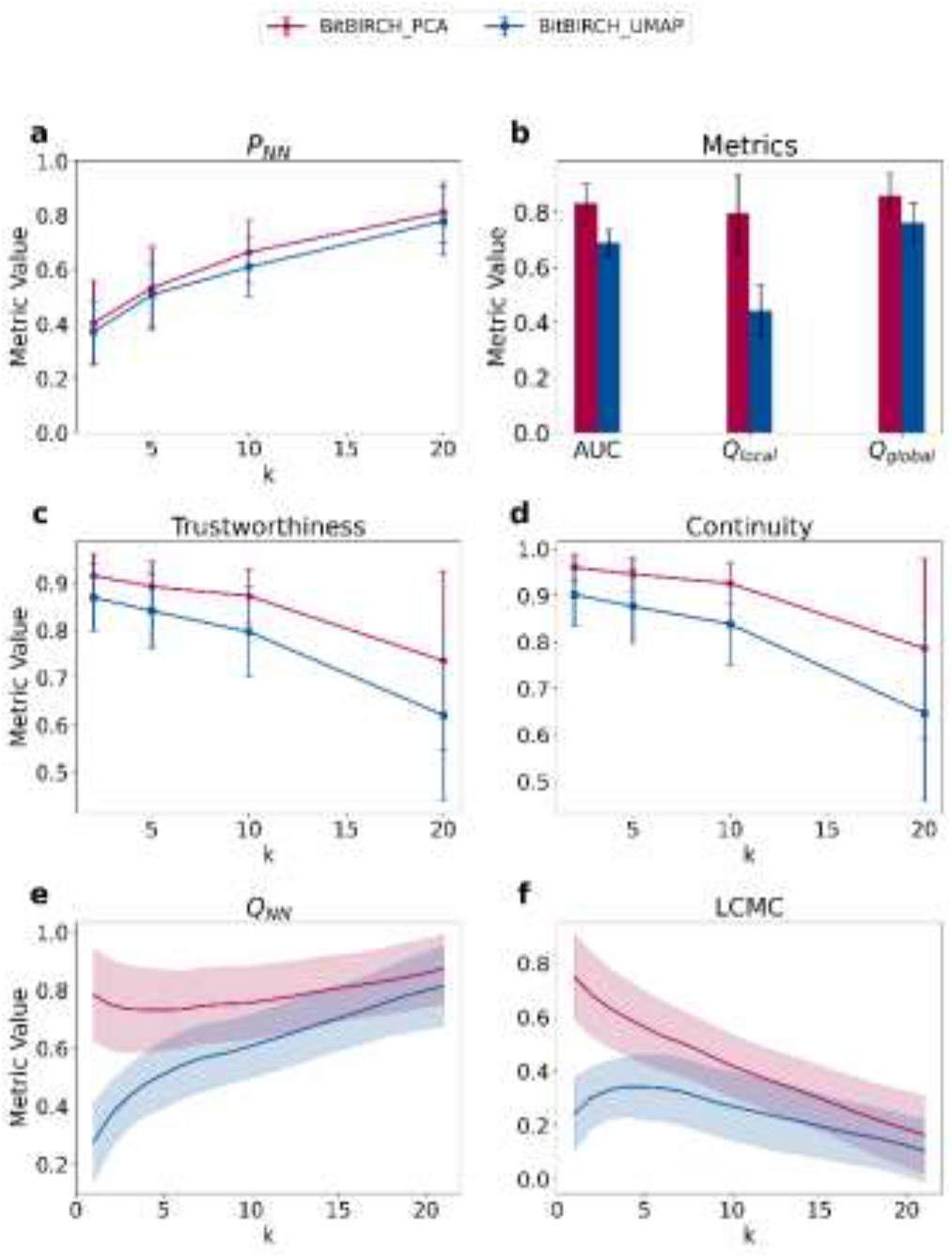
Detailed analysis of neighborhood metrics for the BitBIRCH-then-PCA and BitBIRCH-then-UMAP for clusters having more than 20 molecules.

To test the influence of the initial clustering step, we used BitBIRCH. While there are multiple variants of this algorithm that could also easily be accommodated into this framework, here we decided to use one of the simplest refinement strategies: performing diameter BitBIRCH clustering, with a 0.65 similarity threshold, and then a BF refinement, as described in Ref (63). After the BitBIRCH step, each cluster is projected independently of the others, so essentially we are simultaneously creating maps of different sectors of chemical space. We also considered how these results change if we consider clusters containing more than 10, 20, or 50 molecules, as a way to assess the impact of the degree of granularity. Fig. 8 contains the details of multiple neighborhood metrics averaged over the resulting BitBIRCH clusters, after performing PCA or UMAP. From this, we can draw two immediate key conclusions: all the metrics improved significantly compared to the analysis of the whole set, and PCA now outperforms UMAP (or, at worst, provides equivalent results). Even the AUC metric (Fig. 8A, which was adequate for the whole-set tests) improves from 0.6 to near (and above) 0.8 for both PCA and UMAP. This is also reflected on the *Q*_*global*_ results (Fig. 8B), a case in which we already see a tendency of PCA to outperform UMAP, especially in cases where smaller clusters are considered. The biggest gains (and differences) can be seen in the local metrics. For example, *P*_*NN*_ values (Fig. 8C) are dramatically improved in this new approach, which is perhaps the most important argument in favor of the PAC recipe. Note how in the standard approach PCA values are always below 0.2, while now they are not only always above 0.4, but even get closer to 0.8. Likewise, the *Q*_*local*_ behavior (Fig. 8D) shows a much improved PCA, with this metric now being consistently above 0.6, from a meager 0.3 in the case of the whole set. In these cases, however, the value of *k*_*max*_ is around 1.2 (see SI), so *Q*_*local*_ essentially corresponds to the preservation of the closest nearest neighbor. On the other hand, Trustworthiness and Continuity, which quantify the amount of hard intrusions (distant points that appear close) and hard extrusions (close points that appear distant), provide a fascinating picture. The key difference between *P*_*NN*_ and these metrics is that *P*_*NN*_ is rank-independent and will naturally tend to unity for smaller clusters, as the specific order of neighbors is irrelevant to the metric. In contrast, trustworthiness and continuity are rank-dependent, providing a closer look at the nearest neighbors (beyond just the closest molecule). While these metrics are close for UMAP and BitBIRCH-then-PCA up to *k* = 10, they appear to fall quickly from 10 to 20, also showing large error bars. This was expected since rank dependency of these metrics combined with evaluation over larger neighborhood (e.g., *k* values near the cluster size, here 20) is statistically bound to induce higher error rates. A similar trend was also seen for these metrics in the case of clusters of size of more than 10 and 50 molecules (see SI), providing a strong indication that the decline does not necessarily imply poor performance but the need for a contextual redefinition of what we call the local neighborhood. Moreover, there are significant gains compared to the UMAP results, which are consistent throughout the different granularity levels. These local results are particularly illuminating: since the clustering reduced the size of the neighborhoods to be analyzed, the linear PCA can now capture global and (specially) local information much better, without the burden of trying to find an optimum transformation for the whole set. On the other hand, the reduced (and extremely) local information used to train UMAP diminishes the final projection (mostly an overfitting problem). This is even more remarkable once we note that we are comparing PCA with UMAP after optimizing the hyper-parameters for the latter (that is, these are the best possible UMAP results for these cases), while PCA is, obviously, parameter-free. Add to this that processing all the PCAs for all the clusters only took 30 seconds, while all the UMAPs required 4900 seconds, and the superiority of PCA becomes even more apparent. In short, the BitBIRCH-then-PCA approach is (at worst) equally good as the fine-tuned BitBIRCH-then-UMAP (while being superior in several metrics), but being much more efficient. Moreover, the PAC recipe dramatically outperforms the analysis of the whole set, validating the hypothesis that analyzing local sections of a chemical library provides a much better recipe to map its content to a 2D representation.

**Figure 8.**
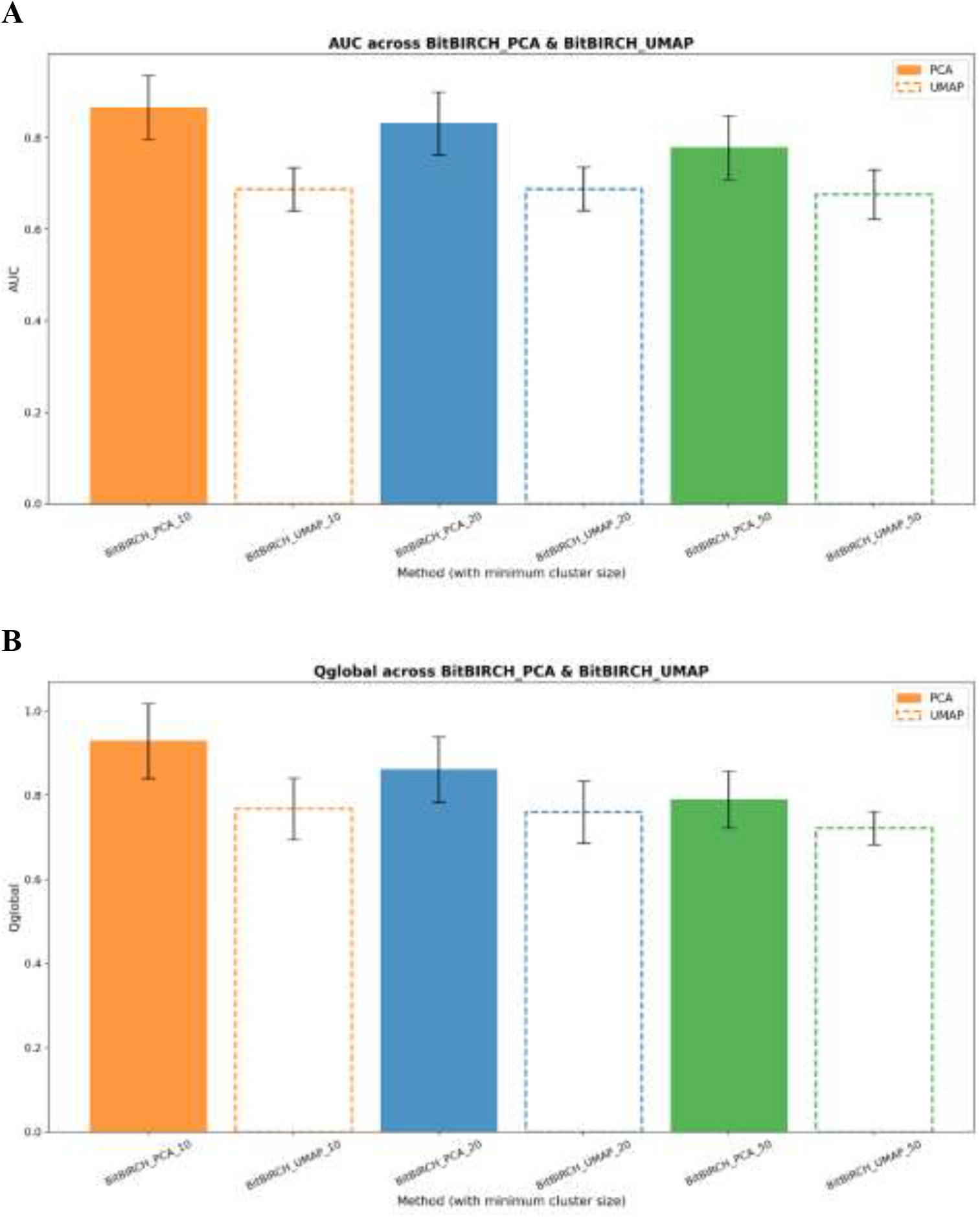
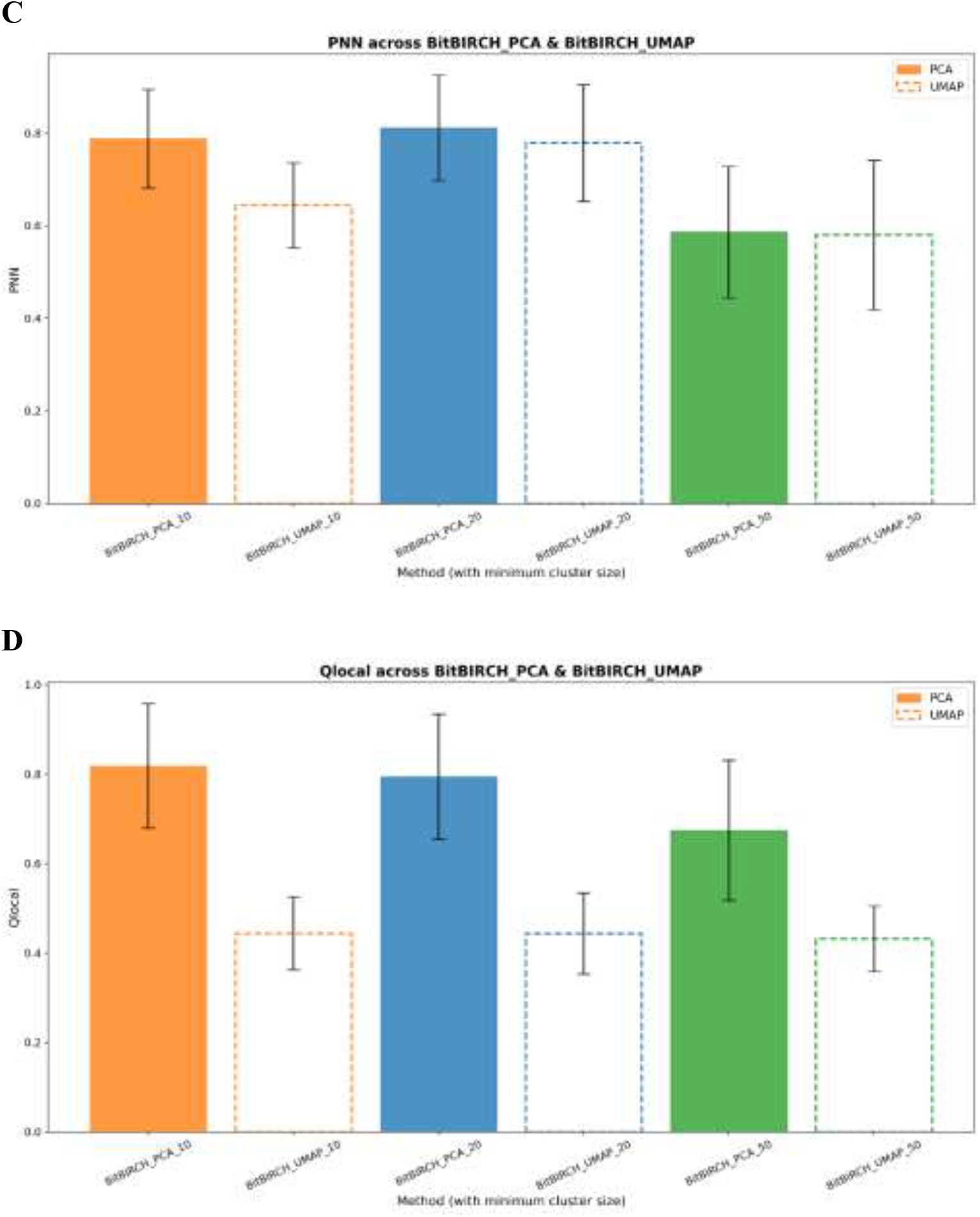
Neighborhood metrics for the project-after-cluster protocol across minimum cluster size of 11, 21 and 51 for CHEMBL33 dataset with ECFP4 fingerprints. PCA (full colors), UMAP (dashed lines). A: AUC, B: *Q*_*global*_, C: *P*_*NN*_, D: *Q*_*local*_.

### 3.4 N-Ary Mapping Interface

With the insights obtained in the previous sections, we then proceeded to build a visualization pipeline, which we implemented as the N-Ary Mapping Interface (NAMI). In short, NAMI provides a two-level view of a given chemical space. First, the user picks a similarity threshold and performs BitBIRCH clustering of the whole set on the original fingerprint space. Note that any flavor of BitBIRCH can be accommodated in this pipeline, including the recently proposed refinement strategies. This step provides the information for the two visualization stages. On one hand, the centroids of the clusters will be used to represent the overall distribution of the high-density molecular regions in a first level of approximation, serving as “markers” for their corresponding clusters. That’s it, we perform the first PCA on the cluster centroids alone. Then, the internal structure of each cluster could be accessed by doing independent PCA projections of the molecules belonging to the cluster, independently of the remaining clusters. At both the centroid level or the cluster level, we have the option to perform the PCA directly over the binary representation of the centroids or fingerprints, or doing the PCA after hyper-plane projection or just the simple normalization.

This implementation was performed in an open-source GUI, which can be accessed in https://github.com/mqcomplab/NAMI. The user has the option to upload just a CSV file containing SMILES and perform the BitBIRCH clustering directly from the NAMI interface (setting the clustering parameters like similarity threshold, branching factor, radius and bits of Morgan fingerprints). However, for convenience of processing large sets, BitBIRCH could be performed independently, and then just loading the final results, including the pre-calculated cluster assignments. To help with the navigation of large sets, we included an option to filter clusters containing a range of molecules as well as to zoom inside a cluster to see the individual molecules and their relative positions in chemical space. Figure 9 shows the centroids for clusters containing more than 250 molecules when we performed the clustering on the ZINC20 dataset containing 10 million molecules. A detailed depiction of a cluster in this dataset is shown in Figure 9B where one can observe the relative position of molecules in the projected two-dimensional PCA space as well as see the structures of the respective molecules. The density within the cluster is calculated using Gaussian kernel density estimation and the lighter regions represent crowded regions. While just a proof-of-principle implementation in its current state, NAMI already provides a faithful representation of chemical space relying only on very efficient tools (BitBIRCH and PCA), by focusing on the locally dense sectors of a library and avoiding the tuning and overfitting of non-linear DR methods.

**Figure 9.**
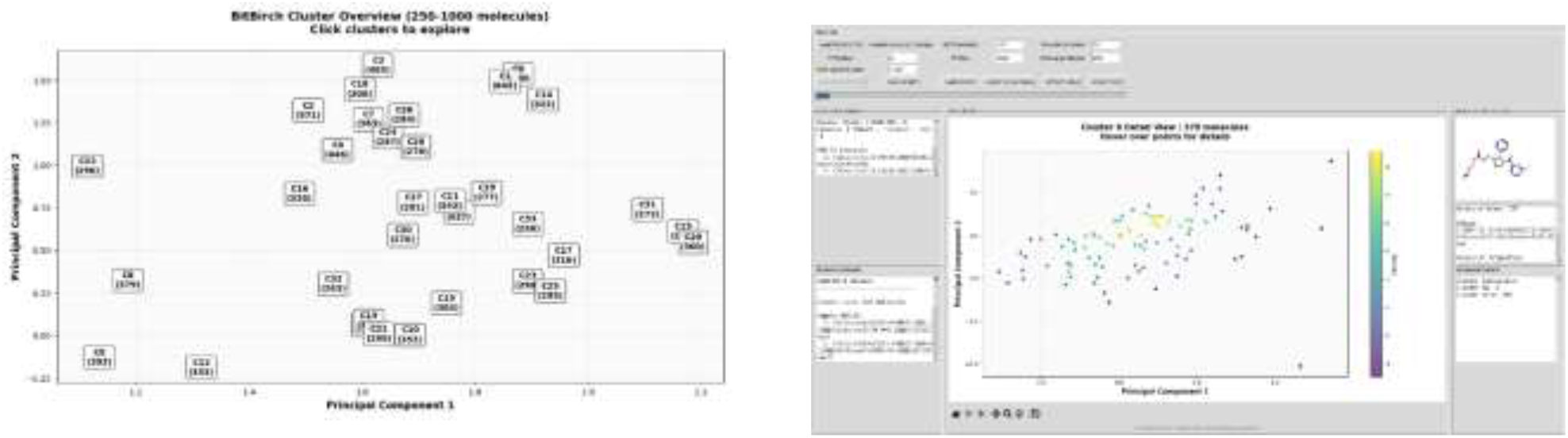
Sample visualization with NAMI of the 10M library selected from ZINC20. Left: Distribution of clusters with more than 250 molecules, projected from the cluster centroids. Right: Local visualization of a cluster.

## 4. CONCLUSIONS

We have shown that linear DR methods can compete with (and even surpass) non-linear alternatives in speed, ease of use, and in neighborhood preservation. The was possible by recognizing that we can exploit the inherent structure of the original high-dimensional fingerprint space. In this regard, the key observation is that the manifold induced by the Tanimoto index in binary fingerprint space can be put in correspondence with a hyper-spherical surface (independently of the chemical library under analysis). This simple relation provides two key insights: the smooth nature of the surface containing the molecules suggests that very flexible dimensionality reduction methods are not needed to capture as much information as possible; and that a more direct implementation of Euclidean-based post-processing tools only requires normalizing the fingerprints at the start of the pipeline. The latter is justified by the excellent ranking correlations between binary Tanimoto and continuous Euclidean distances calculated with normalized representations (in stark contrast with the essentially non-existing correlation with the Euclidean metric calculated over the original fingerprints).

To exploit the newfound structure of the fingerprint manifold, we took inspiration from traditional cartography, by going from the normalized fingerprints, to a hyper-plane projection from the spherical surface, to a final dimensionality reduction to a 2D plane. This simple recipe leads to modest improvements in PCA, which are consistent through different fingerprint types. However, the strategy of performing a sole projection centered around a unique point did not improve the results of the UMAP, t-SNE, and GTM methods, compared to their simple application on the original fingerprints. The main reason for this is that these techniques are already flexible enough to not benefit from the discovered manifold structure. Under these conditions, non-linear DR is still far superior (albeit, after careful selection of the projection hyperparameters), especially for local preservation indicators.

This situation changes, however, when we borrow another cartography idea: performing independent DR after identifying locally dense sectors in the starting fingerprint space. This changes the usual relation between clustering and DR. Traditionally, DR is used as a way to reduce the computational burden of the grouping step (even when the number of dimensions is not the bottleneck in the clustering, and even when this can greatly affect and obscure the final partition). Now, we propose to use the clustering as a way to enhance the performance of the DR, just like when a traveler wants to check the most accurate possible map of a city, it looks for a projection that ignores other parts of the globe. This strategy is possible because now we know we are starting from a smooth, well-behaved, manifold, and while a unique projection can introduce significant distortions, its local sections can be described with simple linear tools. The project-after-clustering (PAC) pipeline improved the PCA results to the point that they are on par with, or even superior to, non-linear DR with optimized hyperparameters. This is a remarkable finding, keeping in mind that this superiority now extends to even local preservation, and was achieved at a small fraction of the computational cost (already over 160 times faster for just 64K molecules), and in a fully black-box way. It is yet again not surprising that non-linear DR does not benefit from the clustering step, since the exposure to reduced training sizes (the individual clusters instead of the whole set) means that they will be more prone to overfitting.

As a way to showcase the power of the PAC recipe, we presented a proof-of-principle package: N-Ary Mapping Interface (NAMI). NAMI uses BitBIRCH in the clustering step, as a way to efficiently process ultra-large sectors of chemical space. This provides a two-level visualization strategy, with a first PCA step performed over the BitBIRCH centroids giving a representation of the global distribution inside the library, and then separate projections for each cluster to dissect their local structure. Even a simple NAMI interface succeeds in promptly visualizing ten million molecules. The minimum computational requirements to perform BitBIRCH + PCA (together with the potential parallelization of the latter over each independent cluster), together with the optimally preserved information in the 2D representation make PAC and NAMI very attractive alternatives in the visualization and analysis of ultra-large sectors of chemical space.

## References

(1) Medina-Franco, J. L.; Martinez-Mayorga, K.; Giulianotti, M. A.; Houghten, R. A.; Pinilla, C. Visualization of the Chemical Space in Drug Discovery. Curr. Comput. Aided Drug Des. 2008, 4 (4), 322–333. 10.2174/157340908786786010.

(2) Cihan Sorkun, M.; Mullaj, D.; Koelman, J. M. V. A.; Er, S. ChemPlot, a Python Library for Chemical Space Visualization. Chemistry–Methods 2022, 2 (7), e202200005. 10.1002/cmtd.202200005.

(3) Reymond, J.-L. The Chemical Space Project. Acc. Chem. Res. 2015, 48 (3), 722–730. 10.1021/ar500432k.

(4) Osolodkin, D. I.; Radchenko, E. V.; Orlov, A. A.; Voronkov, A. E.; Palyulin, V. A.; Zefirov, N. S. Progress in Visual Representations of Chemical Space. Expert Opin. Drug Discov. 2015, 10 (9), 959–973. 10.1517/17460441.2015.1060216.

(5) Scalfani, V. F.; Patel, V. D.; Fernandez, A. M. Visualizing Chemical Space Networks with RDKit and NetworkX. J. Cheminformatics 2022, 14 (1), 87. 10.1186/s13321-022-00664-x.

(6) de la Vega de León, A.; Bajorath, J. Chemical Space Visualization: Transforming Multidimensional Chemical Spaces into Similarity-Based Molecular Networks. Future Med. Chem. 2016, 8 (14), 1769–1778. 10.4155/fmc-2016-0023.

(7) Sosnin, S. Chemical Space Visual Navigation in the Era of Deep Learning and Big Data. Drug Discov. Today 2025, 30 (7), 104392. 10.1016/j.drudis.2025.104392.

(8) Cereto-Massagué, A.; Ojeda, M. J.; Valls, C.; Mulero, M.; Garcia-Vallvé, S.; Pujadas, G. Molecular Fingerprint Similarity Search in Virtual Screening. Methods 2015, 71, 63–. 10.1016/j.ymeth.2014.08.005.

(9) Probst, D.; Reymond, J.-L. A Probabilistic Molecular Fingerprint for Big Data Settings. J. Cheminformatics 2018, 10 (1), 66. 10.1186/s13321-018-0321-8.

(10) Duvenaud, D. K.; Maclaurin, D.; Iparraguirre, J.; Bombarell, R.; Hirzel, T.; Aspuru-Guzik, A.; Adams, R. P. Convolutional Networks on Graphs for Learning Molecular Fingerprints. In Advances in Neural Information Processing Systems; Curran Associates, Inc., 2015; Vol. 28.

(11) Todeschini, R.; Consonni, V. Handbook of Molecular Descriptors; John Wiley & Sons, 2008.

(12) Miranda-Quintana, R.-A.; Cruz-Rodes, R.; Codorniu-Hernandez, E.; Batista-Leyva, A. J. Formal Theory of the Comparative Relations: Its Application to the Study of Quantum Similarity and Dissimilarity Measures and Indices. J. Math. Chem. 2010, 47 (4), 1344–1365. 10.1007/s10910-009-9658-6.

(13) Miranda-Quintana, R. A.; Bajusz, D.; Rácz, A.; Héberger, K. Differential Consistency Analysis: Which Similarity Measures Can Be Applied in Drug Discovery? Mol. Inform. 2021, 40 (7), 2060017. 10.1002/minf.202060017.

(14) Surendran, A.; Zsigmond, K.; López-Pérez, K.; Miranda-Quintana, R. A. Is the Tanimoto Similarity a Metric? J. Math. Chem. 2025, 63 (5), 1229–1240. 10.1007/s10910-025-01721-0.

(15) Kirkpatrick, P.; Ellis, C. Chemical Space. Nature 2004, 432 (7019), 823–824.

(16) Lipinski, C.; Hopkins, A. Navigating Chemical Space for Biology and Medicine. Nature 2004, 432 (7019), 855–861. 10.1038/nature03193.

(17) Reymond, J.-L.; Deursen, R. van; C. Blum L.; Ruddigkeit, L. Chemical Space as a Source for New Drugs. MedChemComm 2010, 1 (1), 30–38. 10.1039/C0MD00020E.

(18) Lopez Perez, K.; López-López, E.; Soulage, F.; Felix, E.; Medina-Franco, J. L.; Miranda-Quintana, R. A. Growth vs Diversity: A Time-Evolution Analysis of the Chemical Space. J. Chem. Inf. Model. 2025, 65 (13), 6788–6796. 10.1021/acs.jcim.5c00347.

(19) Oprea, T. I.; Gottfries, J. Chemography: The Art of Navigating in Chemical Space. J. Comb. Chem. 2001, 3 (2), 157–166. 10.1021/cc0000388.

(20) Gromski, P. S.; Henson, A. B.; Granda, J. M.; Cronin, L. How to Explore Chemical Space Using Algorithms and Automation. Nat. Rev. Chem. 2019, 3 (2), 119–128. 10.1038/s41570-018-0066-y.

(21) Fang, X.; Liu, L.; Lei, J.; He, D.; Zhang, S.; Zhou, J.; Wang, F.; Wu, H.; Wang, H. Geometry-Enhanced Molecular Representation Learning for Property Prediction. Nat. Mach. Intell. 2022, 4 (2), 127–134. 10.1038/s42256-021-00438-4.

(22) Guo, Z.; Guo, K.; Nan, B.; Tian, Y.; Iyer, R. G.; Ma, Y.; Wiest, O.; Zhang, X.; Wang, W.; Zhang, C.; Chawla, N. V. Graph-Based Molecular Representation Learning. arXiv November 29, 2023. 10.48550/arXiv.2207.04869.

(23) Fabian, B.; Edlich, T.; Gaspar, H.; Segler, M.; Meyers, J.; Fiscato, M.; Ahmed, M. Molecular Representation Learning with Language Models and Domain-Relevant Auxiliary Tasks. arXiv November 26, 2020. 10.48550/arXiv.2011.13230.

(24) Harnik, Y.; Milo, A. A Focus on Molecular Representation Learning for the Prediction of Chemical Properties. Chem. Sci. 2024, 15 (14), 5052–5055. 10.1039/D4SC90043J.

(25) Stumpfe, D.; Bajorath, J. Similarity Searching. WIREs Comput. Mol. Sci. 2011, 1 (2), 260–282. 10.1002/wcms.23.

(26) Willett, P. The Calculation of Molecular Structural Similarity: Principles and Practice. Mol. Inform. 2014, 33 (6–7), 403–413. 10.1002/minf.201400024.

(27) The Similarity Principle and Its Application. https://mdpi.org/lin/similarity/similarity-20050529.htm (accessed 2025-09-21).

(28) Maggiora, G. M.; Shanmugasundaram, V. Molecular Similarity Measures. In Chemoinformatics and Computational Chemical Biology; Bajorath, J., Ed.; Humana Press: Totowa, NJ, 2011; pp 39–100. 10.1007/978-1-60761-839-3_2.

(29) Maggiora, G. M. Introduction to Molecular Similarity and Chemical Space; Martinez-Mayorga, K., Medina-Franco, J. L., Eds.; Springer International Publishing: Cham, 2014. 10.1007/978-3-319-10226-9_1.

(30) Medina-Franco, J. L.; Maggiora, G. M. Molecular Similarity Analysis. In Chemoinformatics for Drug Discovery; John Wiley & Sons, Ltd, 2013; pp 343–399. 10.1002/9781118742785.ch15.

(31) Maggiora, G.; Vogt, M.; Stumpfe, D.; Bajorath, J. Molecular Similarity in Medicinal Chemistry. J. Med. Chem. 2014, 57 (8), 3186–3204. 10.1021/jm401411z.

(32) Dunn, T. B.; López-López, E.; Kim, T. D.; Medina-Franco, J. L.; Miranda-Quintana, R. A. Exploring Activity Landscapes with Extended Similarity: Is Tanimoto Enough? Mol. Inform. 2023, 42 (7), 2300056. 10.1002/minf.202300056.

(33) Barbosa, F.; Horvath, D. Molecular Similarity and Property Similarity. Curr. Top. Med. Chem. 2004, 4 (6), 589–600. 10.2174/1568026043451186.

(34) López-Pérez, K.; Miranda-Quintana, R. A. iCliff Taylor’s Version: Robust and Efficient Activity Cliff Determination. J. Chem. Inf. Model. 2025, 65 (11), 5801–5810. 10.1021/acs.jcim.5c00506.

(35) Smale, S. On the Structure of Manifolds. Am. J. Math. 1962, 84 (3), 387–399. 10.2307/2372978.

(36) Munkres, J. R. Analysis On Manifolds; CRC Press: Boca Raton, 2018. 10.1201/9780429494147.

(37) Bishop, R. L.; Crittenden, R. J. Geometry of Manifolds: Geometry of Manifolds; Academic Press, 1964.

(38) Nash, J. C1 Isometric Imbeddings. Ann. Math. 1954, 60 (3), 383–396. 10.2307/1969840.

(39) Nash, J. The Imbedding Problem for Riemannian Manifolds. Ann. Math. 1956, 63 (1), 20–63. 10.2307/1969989.

(40) Greenacre, M.; Groenen, P. J. F.; Hastie, T.; D’Enza, A. I.; Markos, A.; Tuzhilina, E. Principal Component Analysis. Nat. Rev. Methods Primer 2022, 2 (1), 100. 10.1038/s43586-022-00184-w.

(41) Kumar, S.; Nair, A. S.; Bhashkar, V.; Sudevan, S. T.; Koyiparambath, V. P.; Khames, A.; Abdelgawad, M. A.; Mathew, B. Navigating into the Chemical Space of Monoamine Oxidase Inhibitors by Artificial Intelligence and Cheminformatics Approach. ACS Omega 2021, 6 (36), 23399–23411. 10.1021/acsomega.1c03250.

(42) Aouidate, A. Exploring the Chemical Space of BRAF Inhibitors: A Cheminformatic and Machine Learning Analysis. J. Mol. Liq. 2024, 401, 124705. 10.1016/j.molliq.2024.124705.

(43) Choi, J.; Yun, J. S.; Song, H.; Kim, N. H.; Kim, H. S.; Yook, J. I. Exploring the Chemical Space of Protein–Protein Interaction Inhibitors through Machine Learning. Sci. Rep. 2021, 11 (1), 13369. 10.1038/s41598-021-92825-5.

(44) Villares, M.; Saunders, C. M.; Fey, N. Comparison of Dimensionality Reduction Techniques for the Visualisation of Chemical Space in Organometallic Catalysis. Artif. Intell. Chem. 2024, 2 (1), 100055. 10.1016/j.aichem.2024.100055.

(45) Santiago, Á.; Guzmán-Ocampo, D. C.; Aguayo-Ortiz, R.; Dominguez, L. Characterizing the Chemical Space of γ-Secretase Inhibitors and Modulators. ACS Chem. Neurosci. 2021, 12 (15), 2765–2775. 10.1021/acschemneuro.1c00313.

(46) Karlov, D. S.; Sosnin, S.; Tetko, I. V.; Fedorov, M. V. Chemical Space Exploration Guided by Deep Neural Networks. RSC Adv. 2019, 9 (9), 5151–5157. 10.1039/C8RA10182E.

(47) Anders, F.; Chiappini, C.; Santiago, B. X.; Matijevič, G.; Queiroz, A. B.; Steinmetz, M.; Guiglion, G. Dissecting Stellar Chemical Abundance Space with T-SNE. Astron. Astrophys. 2018, 619, A125. 10.1051/0004-6361/201833099.

(48) Andronov, M.; Fedorov, M. V.; Sosnin, S. Exploring Chemical Reaction Space with Reaction Difference Fingerprints and Parametric T-SNE. ACS Omega 2021, 6 (45), 30743–30751. 10.1021/acsomega.1c04778.

(49) Najeeb, J.; Hussain Tahir, M.; Hanafy, A. I.; El-Bahy, S. M.; El-Bahy, Z. M. Machine Learning Assisted Designing of Low Bandgap Indacenodithiophene (IDT)-Based Organic Semi-Conductors. J. Photochem. Photobiol. Chem. 2024, 457, 115877. 10.1016/j.jphotochem.2024.115877.

(50) Olivon, F.; Elie, N.; Grelier, G.; Roussi, F.; Litaudon, M.; Touboul, D. MetGem Software for the Generation of Molecular Networks Based on the T-SNE Algorithm. Anal. Chem. 2018, 90 (23), 13900–13908. 10.1021/acs.analchem.8b03099.

(51) McInnes, L.; Healy, J.; Melville, J. UMAP: Uniform Manifold Approximation and Projection for Dimension Reduction. arXiv September 18, 2020. 10.48550/arXiv.1802.03426.

(52) Ehiro, T. Feature Importance-Based Interpretation of UMAP-Visualized Polymer Space. Mol. Inform. 2023, 42 (8–9), 2300061. 10.1002/minf.202300061.

(53) Joswiak, M.; Peng, Y.; Castillo, I.; Chiang, L. H. Dimensionality Reduction for Visualizing Industrial Chemical Process Data. Control Eng. Pract. 2019, 93, 104189. 10.1016/j.conengprac.2019.104189.

(54) Zabolotna, Y.; Bonachera, F.; Horvath, D.; Lin, A.; Marcou, G.; Klimchuk, O.; Varnek, A. Chemspace Atlas: Multiscale Chemography of Ultralarge Libraries for Drug Discovery. J. Chem. Inf. Model. 2022, 62 (18), 4537–4548. 10.1021/acs.jcim.2c00509.

(55) Bishop, C. M.; Svensén, M.; Williams, C. K. I. GTM: The Generative Topographic Mapping. Neural Comput. 1998, 10 (1), 215–234. 10.1162/089976698300017953.

(56) Gaspar, H. A.; Baskin, I. I.; Marcou, G.; Horvath, D.; Varnek, A. GTM-Based QSAR Models and Their Applicability Domains. Mol. Inform. 2015, 34 (6–7), 348–356. 10.1002/minf.201400153.

(57) Sidorov, P.; Gaspar, H.; Marcou, G.; Varnek, A.; Horvath, D. Mappability of Drug-like Space: Towards a Polypharmacologically Competent Map of Drug-Relevant Compounds. J. Comput. Aided Mol. Des. 2015, 29 (12), 1087–1108. 10.1007/s10822-015-9882-z.

(58) Zabolotna, Y.; Ertl, P.; Horvath, D.; Bonachera, F.; Marcou, G.; Varnek, A. NP Navigator: A New Look at the Natural Product Chemical Space. Mol. Inform. 2021, 40 (9), 2100068. 10.1002/minf.202100068.

(59) Janssen, A. P. A.; Grimm, S. H.; Wijdeven, R. H. M.; Lenselink, E. B.; Neefjes, J.; van Boeckel, C. A. A.; van Westen, G. J. P.; van der Stelt, M. Drug Discovery Maps, a Machine Learning Model That Visualizes and Predicts Kinome–Inhibitor Interaction Landscapes. J. Chem. Inf. Model. 2019, 59 (3), 1221–1229. 10.1021/acs.jcim.8b00640.

(60) Orlov, A. A.; Akhmetshin, T. N.; Horvath, D.; Marcou, G.; Varnek, A. From High Dimensions to Human Insight: Exploring Dimensionality Reduction for Chemical Space Visualization. Mol. Inform. 2025, 44 (1), e202400265. 10.1002/minf.202400265.

(61) Surendran, A.; Zsigmond, K.; Miranda-Quintana, R. A. Undersampling Techniques for Non-Linear Chemical Space Visualization. bioRxiv 2025, 2025.07.03.663077. 10.1101/2025.07.03.663077.

(62) Pérez, K. L.; Jung, V.; Chen, L.; Huddleston, K.; Miranda-Quintana, R. A. BitBIRCH: Efficient Clustering of Large Molecular Libraries. Digit. Discov. 2025, 4 (4), 1042–1051. 10.1039/D5DD00030K.

(63) López Pérez, K.; Huddleston, K.; Jung, V.; Miranda-Quintana, R. A. BitBIRCH Clustering Refinement Strategies. J. Chem. Inf. Model. 2025, 65 (11), 5280–5288. 10.1021/acs.jcim.5c00627.

(64) Lopez-Perez, K.; Zhao, B.; Miranda-Quintana, R. A. iSIM-Sigma: Efficient Standard Deviation Calculation for Molecular Similarity. J. Chem. Inf. Model. 2025, 65 (13), 6797–6808. 10.1021/acs.jcim.5c00894.

(65) López-Pérez, K.; Kim, T. D.; Miranda-Quintana, R. A. iSIM: Instant Similarity. Digit. Discov. 2024, 3 (6), 1160–1171. 10.1039/D4DD00041B.

(66) Miranda-Quintana, R. A.; Bajusz, D.; Rácz, A.; Héberger, K. Extended Similarity Indices: The Benefits of Comparing More than Two Objects Simultaneously. Part 1: Theory and Characteristics†. J. Cheminformatics 2021, 13 (1), 32. 10.1186/s13321-021-00505-3.

(67) Miranda-Quintana, R. A.; Rácz, A.; Bajusz, D.; Héberger, K. Extended Similarity Indices: The Benefits of Comparing More than Two Objects Simultaneously. Part 2: Speed, Consistency, Diversity Selection. J. Cheminformatics 2021, 13 (1), 33. 10.1186/s13321-021-00504-4.

(68) Besau, F.; Hack, T.; Pivovarov, P.; Schuster, F. E. Spherical Centroid Bodies. Am. J. Math. 2023, 145 (2), 515–542. 10.1353/ajm.2023.0012.

(69) nLab authors. stereographic projection. https://ncatlab.org/nlab/show/stereographic+projection.

(70) van Tilborg, D.; Alenicheva, A.; Grisoni, F. Exposing the Limitations of Molecular Machine Learning with Activity Cliffs. J. Chem. Inf. Model. 2022, 62 (23), 5938–5951. 10.1021/acs.jcim.2c01073.

(71) Zdrazil, B.; Felix, E.; Hunter, F.; Manners, E. J.; Blackshaw, J.; Corbett, S.; de Veij, M.; Ioannidis, H.; Lopez, D. M.; Mosquera, J. F.; Magarinos, M. P.; Bosc, N.; Arcila, R.; Kizilören, T.; Gaulton, A.; Bento, A. P.; Adasme, M. F.; Monecke, P.; Landrum, G. A.; Leach, A. R. The ChEMBL Database in 2023: A Drug Discovery Platform Spanning Multiple Bioactivity Data Types and Time Periods. Nucleic Acids Res. 2024, 52 (D1), D1180–D1192. 10.1093/nar/gkad1004.

(72) CHEMBL33. 10.6019/CHEMBL.database.33.

(73) Rogers, D.; Hahn, M. Extended-Connectivity Fingerprints. J. Chem. Inf. Model. 2010, 50 (5), 742–754. 10.1021/ci100050t.

(74) Irwin, J. J.; Tang, K. G.; Young, J.; Dandarchuluun, C.; Wong, B. R.; Khurelbaatar, M.; Moroz, Y. S.; Mayfield, J.; Sayle, R. A. ZINC20—A Free Ultralarge-Scale Chemical Database for Ligand Discovery. J. Chem. Inf. Model. 2020, 60 (12), 6065–6073. 10.1021/acs.jcim.0c00675.

(75) Rogers, D. J.; Tanimoto, T. T. A Computer Program for Classifying Plants. Science 1960, 132 (3434), 1115–1118. 10.1126/science.132.3434.1115.

(76) Kendall, M. G. The Treatment of Ties in Ranking Problems. Biometrika 1945, 33, 251–. 10.1093/biomet/33.3.239.

