## Supplementary Information for "The Shape of Chemical Space"

### S1. BitBIRCH then PCA/UMAP for ECFP4 fingerprints

Complete analysis for clusters containing more than 10 and 50 molecules.

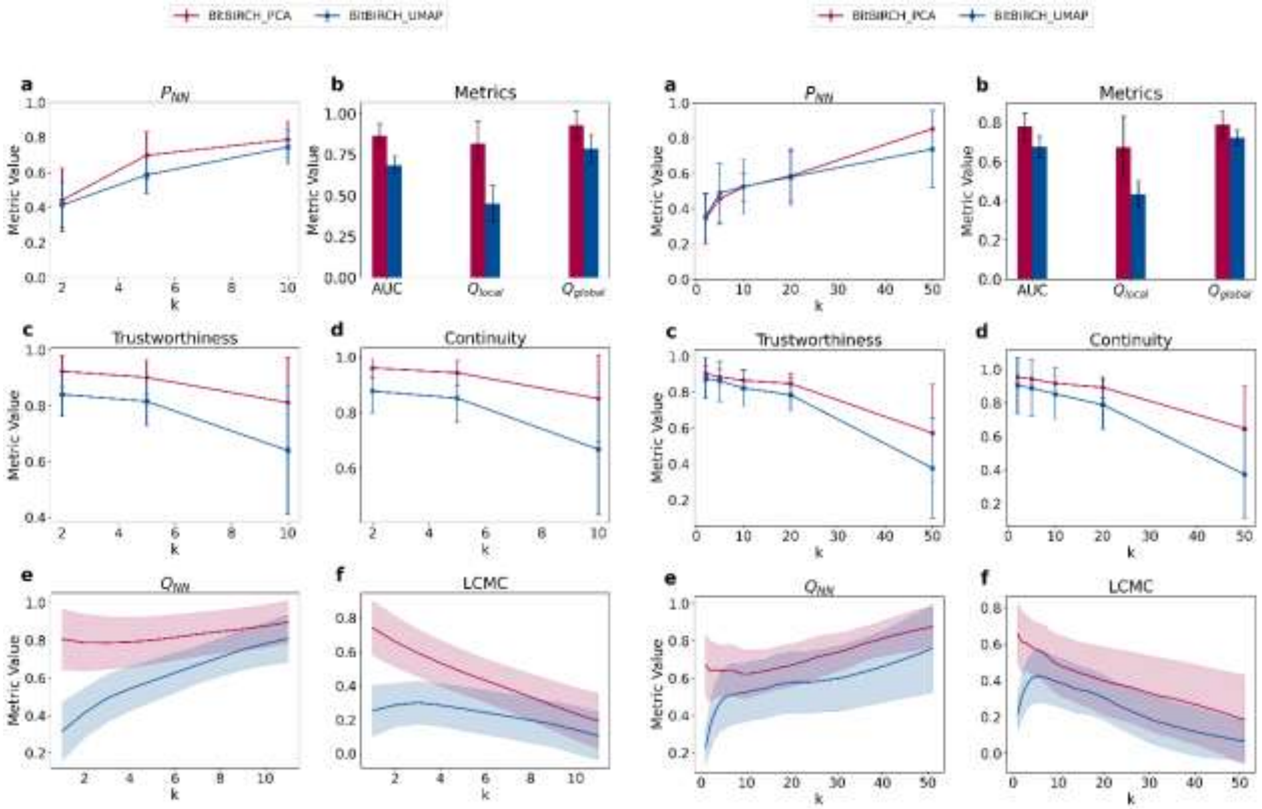


**Figure S1:** Detailed analysis of neighborhood metrics for the BitBIRCH-then-PCA and BitBIRCH-then-UMAP for ECFP4 fingerprint clusters having more than left: 10 and right: 50 molecules.

Figure S1 shows the results for neighborhood preservation metrics for the BitBIRCH-then-PCA and BitBIRCH-then-UMAP paradigms applied to ECFP fingerprint clusters. Consistent with the results for clusters with more than 20 molecules in the main text, the local neighborhood preservation metric $P_{NN}$ remains nearly identical between both approaches, while rank based metrics like Trustworthiness and Continuity in BitBIRCH-then-PCA slightly overperform those in BitBIRCH-then-UMAP. Notably, a steeper decline is observed for these metrics at higher neighborhood sizes (large k), indicating higher rank discrepancies when broader neighborhoods are considered. The results are consistent across all minimum cluster sizes, and the following section deals with how the choice of an alternative fingerprint representation could potentially influence the quality of dimensionality reduction.

**S2. Comparison of dimensionality reduction metrics for RDKit fingerprints**

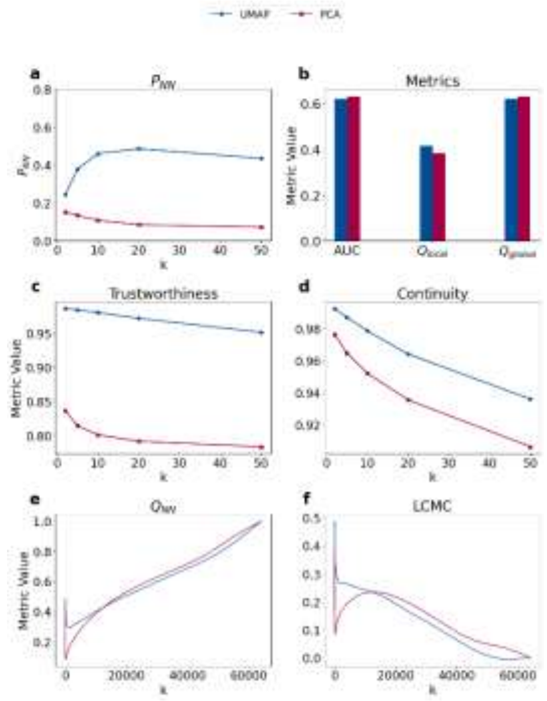


**Figure S2:** Detailed analysis of neighborhood metrics for the 64K set after projecting using PCA and UMAP for the 2048-dimensional RDKit fingerprints.

An analogous analysis to Figure 6 of the main text, now utilizing 2048-dimensional RDKit fingerprints instead of ECFP4 shows little to no differences to the results of ECFP, indicating that these dimensionality reduction methods perform consistently regardless of the dimensionality of the fingerprint vector (1024 or 2048). As expected, local neighborhood preservation metrics like $P_{NN}$, Trustworthiness and Continuity portray a superior performance for UMAP to PCA.

We performed the cluster-then-project procedure for RDKit fingerprints to check for consistency in the improvement of projection metrics across molecular representations. Figure S3 presents the results for clusters containing a minimum of 11, 21 and 51 molecules, respectively. As before, we performed diameter BitBIRCH with a 0.65 similarity threshold and BF refinement. The results are again consistent to the one in the main text, with BitBIRCH-PCA consistently delivering reliable projections for yet another fingerprint type. Local metrics like $P_{NN}$, Trustworthiness and Continuity show better performance across all k values for all cluster size settings for BitBIRCH-PCA, aligning with the trends observed for ECFP4 fingerprints.

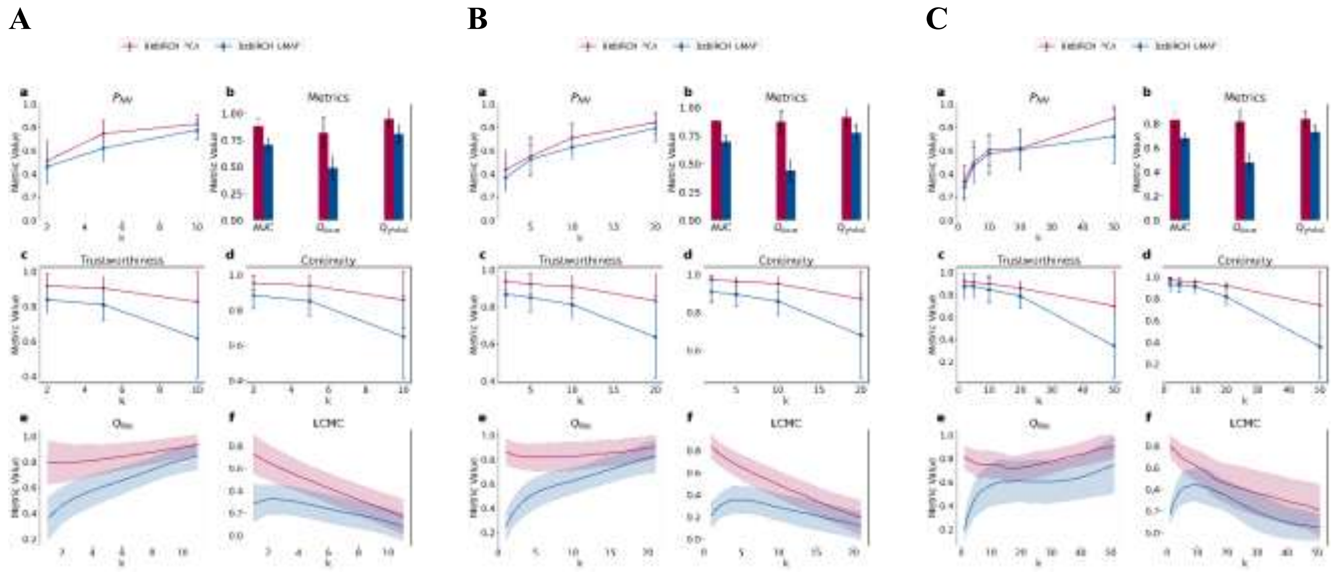


**Figure S3:** Detailed analysis of neighborhood metrics for the BitBIRCH-then-PCA and BitBIRCH-then-UMAP on RDKit fingerprints for clusters having at least A) 11, B) 21 and C) 51 molecules.

#### S3. Comparison of DR metrics across Normalized and Gnomonic centroid projections

Building upon the cartography results for PCA, we attempt to see if performing these projections before the BitBIRCH-then-PCA recipe provides any profound insights. The results for BitBIRCH-PCA performed on Normalized fingerprints and Gnomonic centroid projected fingerprints are shown in Figure S4 and S5 respectively. The metric values and trends appear closely similar to those in the plain ECFP4 case, indicating that projecting prior to dimensionality reduction offers little to no change in the quality of neighborhood preservation.

A B C

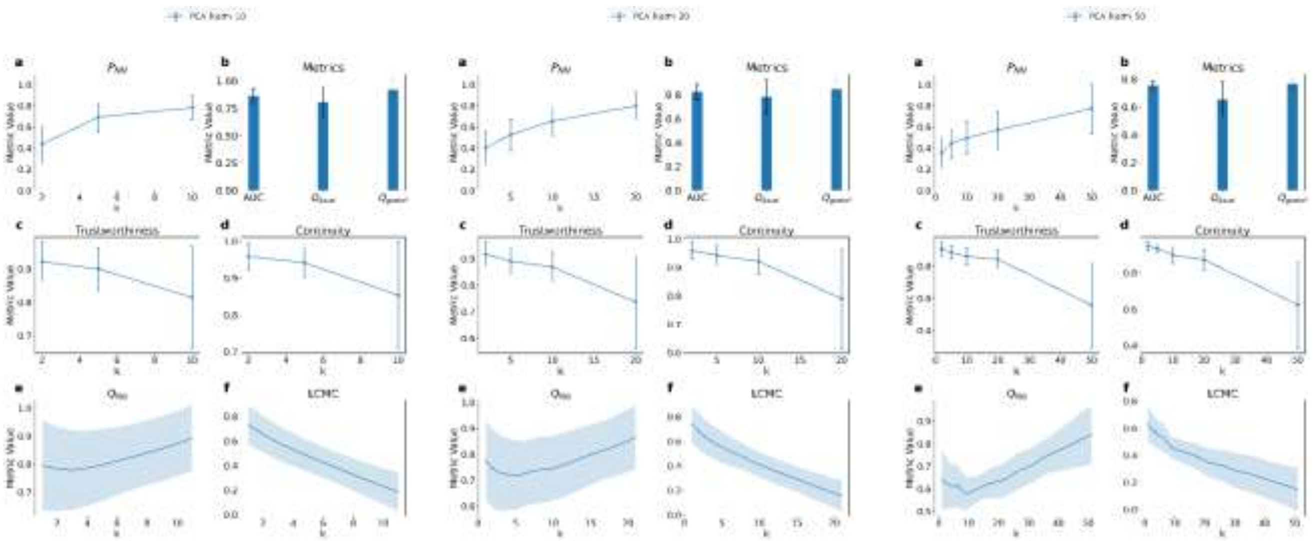


**Figure S4:** Detailed analysis of neighborhood metrics for the BitBIRCH-then-PCA on Normalized ECFP4 fingerprints for clusters having at least A) 11, B) 21 and C) 51 molecules.

A B C

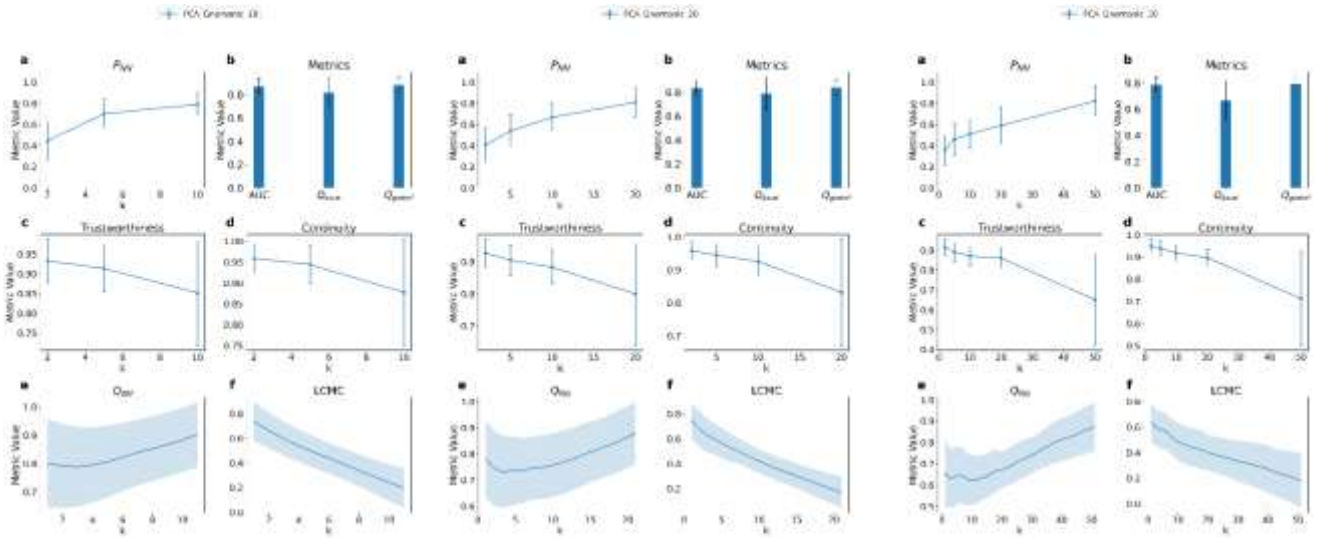

**Figure S5:** Detailed analysis of neighborhood metrics for the BitBIRCH-then-PCA on Gnomonic centroid projected fingerprints for clusters having at least A) 11, B) 21 and C) 51 molecules.

### S4. A closer look at the distribution of metrics across clusters

The observed trend in neighborhood metrics across different molecular representations and projection methods affirm the robustness of BitBIRCH-PCA pipeline for chemical space visualization. This linear approach consistently yields lower information loss compared to non-linear alternatives like UMAP, providing a fast and reliable option, especially for large chemical datasets and scenarios where hyperparameter tuning poses practical challenges. To further refine these insights, a final granular analysis was performed by examining the distribution of neighborhood preservation metrics at the individual cluster level.

Figure S6 illustrates these distributions across all the clusters (ordered in decreasing order of cluster size). Out of all the metrics, AUC most clearly differentiates the two methods, with BitBIRCH-PCA exhibiting consistently superior performance for most clusters. On the other hand, local metrics like $P_{NN}$, Trustworthiness and Continuity reveal a more nuanced picture: for the largest clusters, both methods perform comparably, but BitBIRCH-PCA increasingly outperforms BitBIRCH-UMAP as cluster size decreases. Collectively, the evidence positions BitBIRCH-PCA as a robust, linear alternative matching UMAP's (best hyperparameters) projection quality for larger clusters while surpassing it for smaller ones, all while offering significant practical advantages-speed, reproducibility and lack of dataset-specific hyperparameter dependence, making it a suitable choice for large-scale chemical data visualization.

**A**

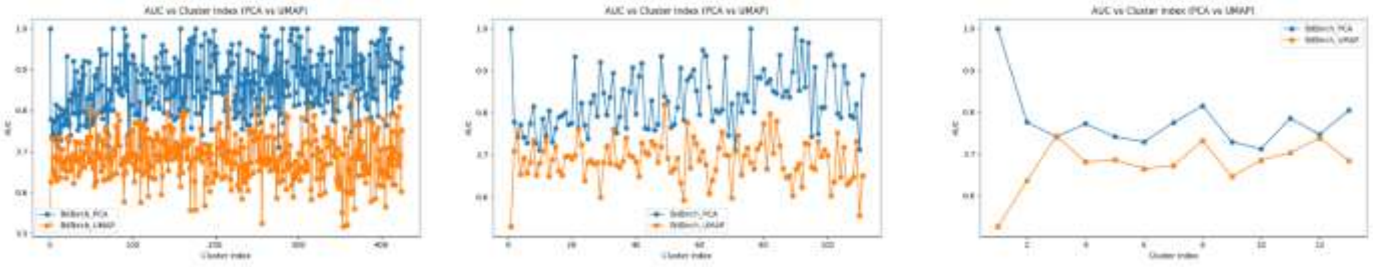

**B**

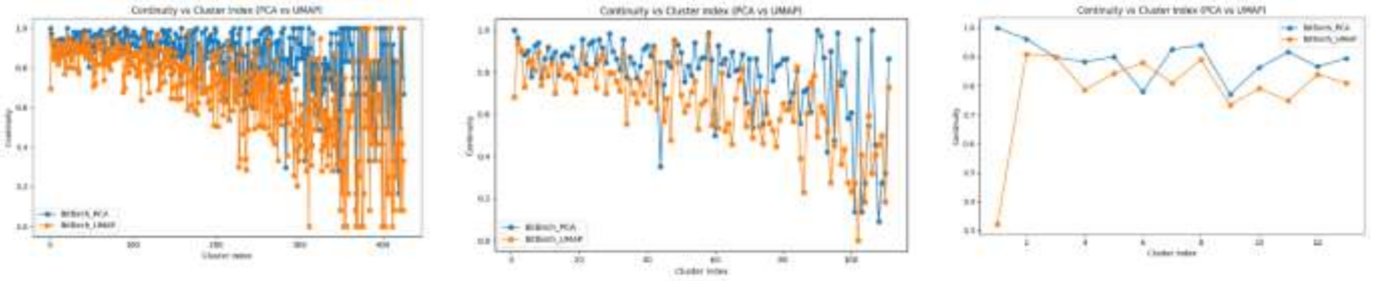


**C**

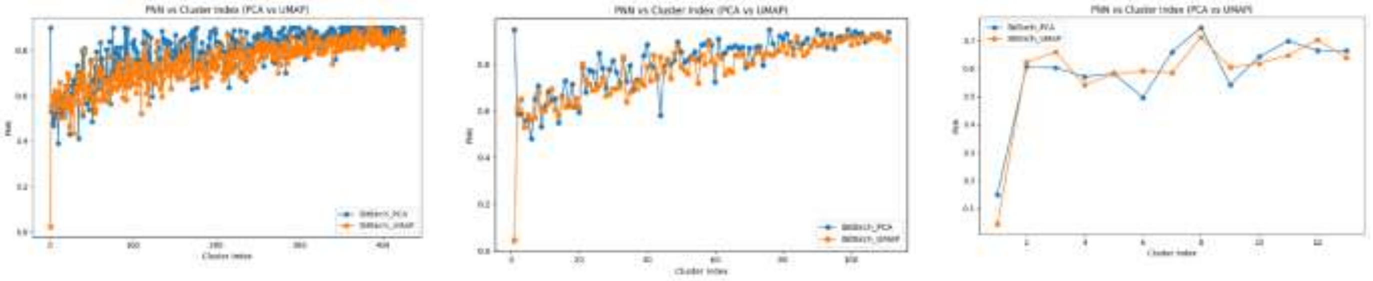


**D**

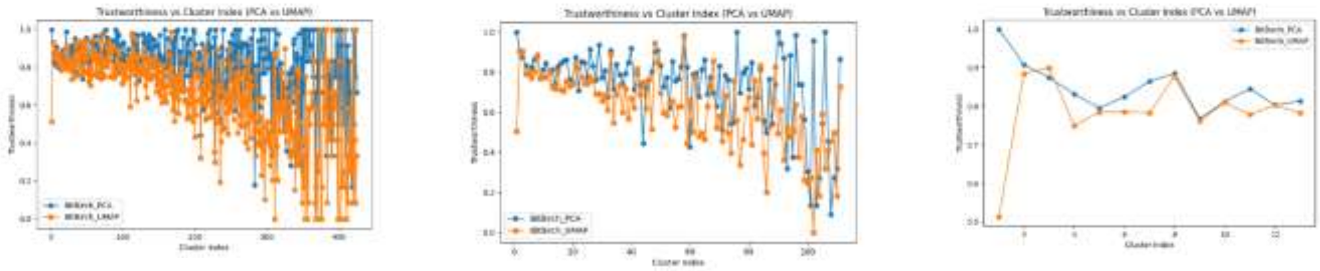


**Figure S6:** Variation of DR metrics: A) AUC, B) Continuity, C) PNN and D) Trustworthiness across individual clusters for BitBIRCH-PCA (blue) and BitBIRCH-UMAP (orange). Clusters indices are arranged in decreasing order of cluster population.
